# The lipid peroxidation product 4-hydroxynonenal inhibits NLRP3 inflammasome activation and macrophage pyroptosis

**DOI:** 10.1101/2022.02.01.478693

**Authors:** Chia George Hsu, Camila Lage Chávez, Chongyang Zhang, Mark Sowden, Chen Yan, Bradford C. Berk

**Affiliations:** Department of Medicine, Aab Cardiovascular Research Institute, University of Rochester School of Medicine and Dentistry, Rochester, NY, USA

**Keywords:** Pyroptosis, 4-hydroxynonenal, Macrophage, NLRP3, Inflammasome, NEK7

## Abstract

Pyroptosis is a form of cell death triggered by the innate immune system that has been implicated in the pathogenesis of sepsis and acute lung injury. At the cellular level, pyroptosis is characterized by cell swelling, membrane rupture, and release of inflammatory cytokines, such as IL-1β. However, the role of endogenous lipids in pyroptosis remains underappreciated. We discovered that 4-hydroxynonenal (HNE), a major endogenous product of lipid peroxidation, inhibited pyroptosis and inflammasome activation. HNE at physiological concentrations (3 µM) blocked nigericin and ATP-induced cell death, as well as secretion of IL-1β, by mouse primary macrophages and human peripheral blood mononuclear cells. Treatment with HNE, or an increase of endogenous HNE by inhibiting glutathione peroxidase 4, reduced inflammasome activation in mouse models of acute lung injury and sepsis. Mechanistically, HNE inhibited the NLRP3 inflammasome activation independently of Nrf2 and NF-κB signaling, and had no effect on the AIM2 inflammasome. Furthermore, HNE directly bound to NLRP3 and inhibited its interaction with NEK7. Our findings identify HNE as a novel, endogenous inhibitor of the NLRP3 inflammasome.

## Introduction

Pyroptosis is a lytic form of programmed cell death, characterized by cytoplasmic swelling, pore formation in the cell membrane, and release of pro-inflammatory cytokines (1). Pyroptosis is initiated in response to pathogen-associated molecular patterns (PAMPs) and damage-associated molecular patterns (DAMPs) via pattern recognition receptors (PRRs) (2). Based on their location, PRRs are divided into membrane-bound PRRs and cytoplasmic PRRs. Toll-like receptors (TLRs) are transmembrane proteins that play important roles in the innate immune response (3). For example, activation of the TLR4 receptor by gram-negative bacteria endotoxins, such as lipopolysaccharide (LPS), stimulates multiple signaling pathways in macrophages, including NF-κB, and the subsequent production of pro-inflammatory cytokines (3). Unlike TLRs, the nucleotide-binding oligomerization domain-like receptors (NOD-like receptors, NLRs) recognize endogenous danger or stress responses, and form multiprotein complexes called inflammasomes (4-6). NLR Family Pyrin Domain Containing 3 (NLRP3), NLR family CARD domain-containing protein 4 (NLRC4) and Absent In Melanoma 2 (AIM2) are the best characterized inflammasomes, and have been implicated in the pathogenesis of sepsis, atherosclerosis, and acute lung injury (4-7). Stimulation of inflammasomes involves two signals: 1) transcriptional and posttranslational priming of inflammasome components, for example by LPS ; and 2) activation of inflammasome assembly by a cell danger signal, such as K^+^ influx, extracellular ATP for NLRP3, or double-stranded DNA for AIM2. The formation of inflammasomes triggers the activation of caspase-1 and subsequent processing of interleukin-1β (IL-1β) and interleukin-18 (IL-18) into their mature forms (8, 9). Gasdermin-D (GSDMD) was discovered as a pore-forming protein and the final effector downstream of caspase-1 activation. Active caspases cleave GSDMD to generate an N-terminal cleavage product (GSDMD-NT) that forms transmembrane pores to enable IL-1β release and to drive pyroptosis (10-12). The importance of IL-1β as a disease mediator was confirmed by the CANTOS trial in which an IL-1β neutralizing antibody led to a lower rate of recurrent cardiovascular events in patients with previous myocardial infarction (13).

Recent data suggest that endogenous lipids or their oxidation products can activate or inhibit the assembly of inflammasomes (5, 14). Among reactive aldehydes derived from lipid peroxidation, 4-hydroxynonenal (HNE) is the most abundant end-product. The concentration of HNE in human serum is 0.05-0.15 μM under physiological conditions (15). However, HNE levels may reach 3-6 μM in tissues under oxidative stress (16, 17). Because of its high solubility in aqueous fluids, the reactive HNE formed in membranes can diffuse into the cytoplasm. HNE is detoxified by conjugation to glutathione by glutathione S-transferase (18, 19). However, some HNE molecules escape this mechanism and react with the side chains of cysteine, histidine and lysine residues in proteins (20-22). HNE thus has emerged as an important second messenger signaling molecule (18, 19). For example, low concentrations of HNE produce beneficial effects, including the stimulation of endogenous antioxidant defense mechanisms and the inhibition of inflammation (19, 23-26). Currently, two mechanisms are proposed for HNE-mediated regulation of inflammation. 1) HNE facilitates antioxidant expression by activating Nrf2 signaling, via disrupting Keap1−Nrf2 association and preventing Nrf 2 degradation (24, 25, 27). Nrf2 stimulates antioxidant expression and increases the resistance to cytotoxic reactive oxygen species (ROS), thereby blocking multiple inflammatory pathways. 2). HNE blocks NF-κB activation by inhibiting IκB kinase (IKK) activity, likely by covalently modifying cysteine residue(s) of IKK (26, 28). In this study we explored a novel mechanism by which the lipid peroxidation product HNE inhibits the NLRP3 inflammasome by directly disrupting the binding of NEK7 to NLRP3, thereby preventing activation of caspase-1. We demonstrate the importance of this pathway in decreasing inflammatory cytokine release and macrophage pyroptosis in vitro and in vivo.

## Results

### HNE inhibits pyroptotic cell death in human and mouse macrophages

Nigericin is a K^+^ ionophore that activates the NLRP3 inflammasome (29) and stimulates pyroptotic cell death. To study the role of HNE in pyroptosis, we used LPS co-incubated with HNE, followed by nigericin, in differentiated human THP-1 macrophages and mouse bone-marrow-derived macrophages (BMDMs). Cell death was confirmed by morphological changes, lactate dehydrogenase (LDH) release, and real-time nucleic acid staining (SYTOX^TM^). HNE treatment alone (3 µM) had no effect on cell death, but HNE significantly decreased the magnitude of LPS-nigericin stimulated cell death as indicated by membrane blebbing, LDH release, and nucleic acid staining in mouse and human macrophages (Fig. 1A-F). These results show that HNE protects macrophages from LPS/nigericin-mediated pyroptosis.

**Fig. 1.**
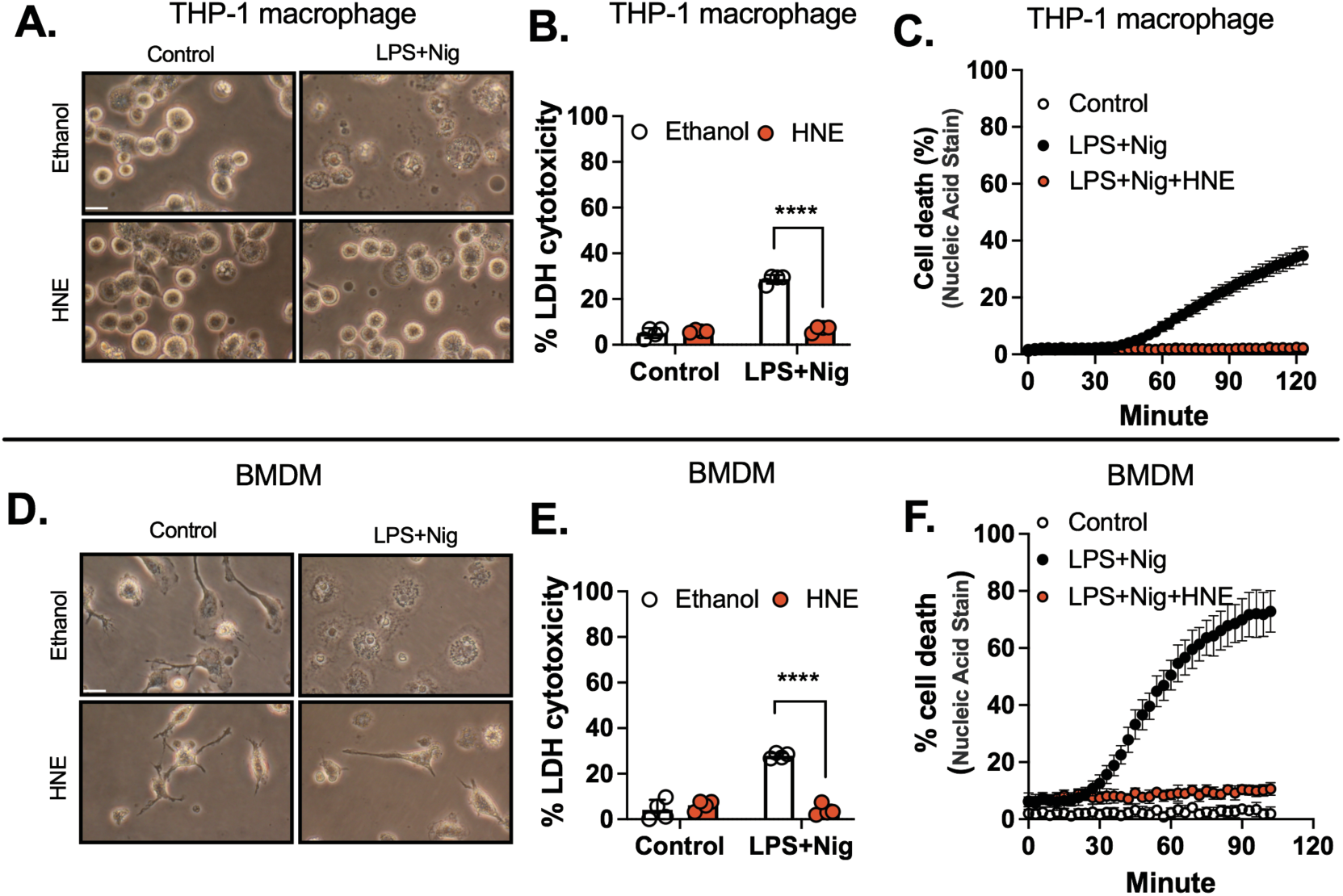
HNE inhibits pyroptotic cell death in human and mouse macrophages. **A-C**. THP-1 differentiated macrophages were stimulated with LPS (100 ng/mL) and co-incubated with HNE (3 μM) or vehicle (ethanol) for 3 hr followed by 2 hr of 6 μM nigericin (Nig) treatment. (A) Cell morphology, scale bar = 10 µm. (B) LDH cytotoxicity. (C) Cell death by SYTOX^TM^ green. **D-F.** Bone-marrow-derived macrophages (BMDMs) were stimulated with LPS (100 ng/mL) and co-incubated with HNE (3 μM) or vehicle (ethanol) for 3 hr followed by 1 hr of 2 μM nigericin (Nig) treatment. (D) Cell morphology, scale bar = 10 µm. (E) LDH cytotoxicity. (F) Cell death by SYTOX^TM^ green. Statistics in B and E were performed using a 2-way ANOVA and Bonferroni’s post hoc test. ****P<0.001 between LPS+Nig+ethanol and LPS+Nig+HNE groups. (N=4 experiments). Bars represent mean ± SD.

### HNE inhibits pyroptosis independent of Nrf2 signaling

Nrf2, an antioxidant transcription factor, has been proposed to regulate NLRP3 inflammasome activation (30-32). Nrf2 function is inhibited by Keap1, which binds Nrf2, facilitates its degradation, and prevents its nuclear translocation (33). To study the potential role of Nrf2 in HNE inhibition of pyroptosis, we first showed that HNE (3 µM) induced Nrf2 activation in macrophages as indicated by its nuclear translocation (Fig. 2A-B), wherase LPS did not. We next studied the effect of HNE on Nrf2 regulated genes, such as glutamate-cysteine ligase catalytic subunit (GCLC) and ferroportin-1 (Fig. 2C-D) (34). LPS treatment alone had no significant effect on any of these measurements (Fig. 2C-D). Simultaneous treatment with LPS and HNE significantly increased Nrf2 activation, compared to LPS alone (Fig. 2A-D). To determine if Nrf2 activation was required for HNE inhibition of pyroptosis, we blocked Nrf2 signaling using the specific inhibitor, ML385 (35) (Fig. S1A-B). We found that 3 µM HNE still prevented cell death in the presence of ML385 (Fig. 2E-F), suggesting that the protective effect of HNE against pyroptosis was independent of the Nrf2 pathway.

**Fig. 2.**
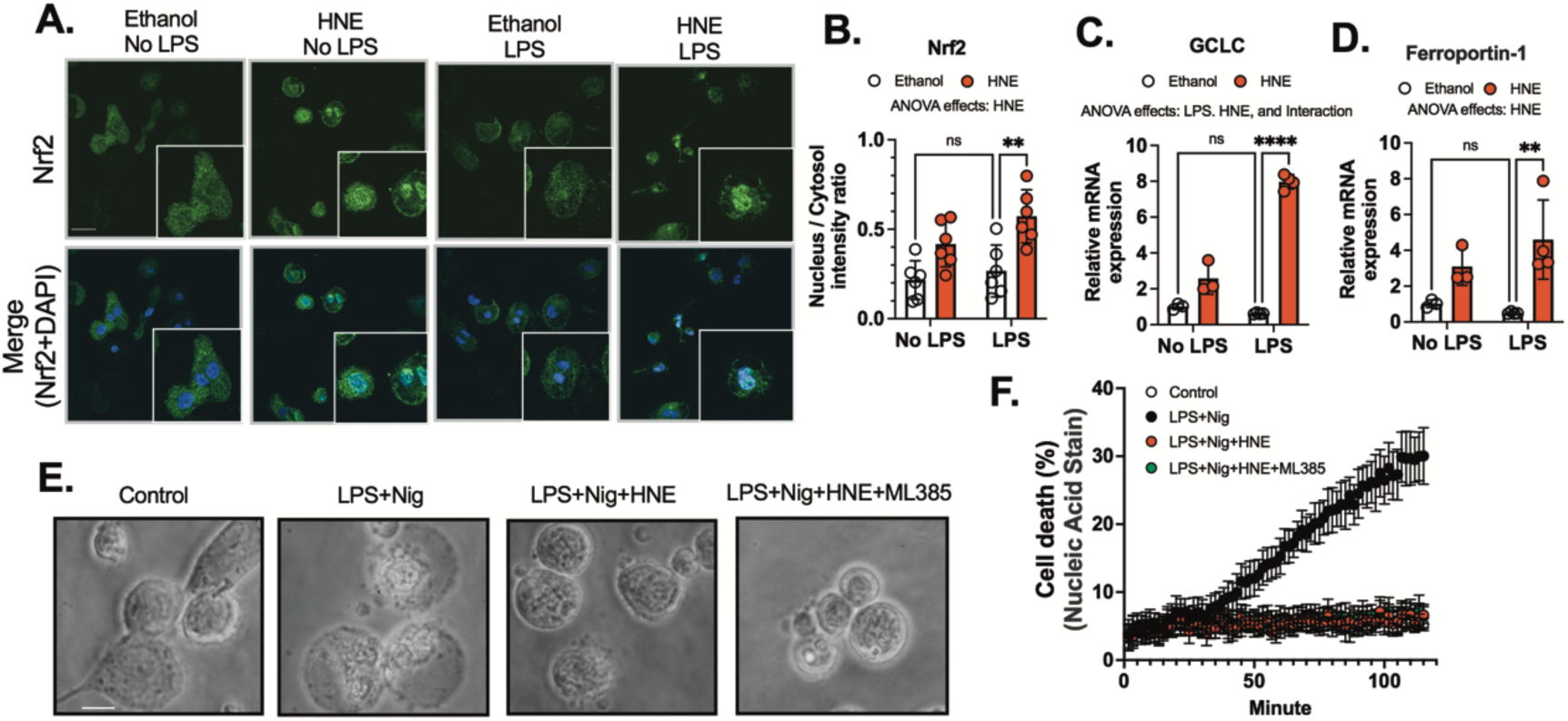
HNE inhibits pyroptosis independent of Nrf2 signaling. **A-B**. THP-1 macrophages were stimulated with or without LPS (100 ng/mL) and co-incubated with ethanol or 3 µM HNE for 3 hr. (A) Nrf2 translocation was determined by immunofluorescence (scale bar: 10 µm), DAPI (blue), and Nrf2 (Green). (B) Nuclear to cytosolic Nrf2 intensity ratio was quantified by Image J. **C-D.** Peritoneal macrophages were stimulated with or without LPS (100 ng/mL) and co-incubated with ethanol or 3 µM HNE for 3 hr. Gene expression was analyzed by real-time PCR. (C) GCLC mRNA expression and (D) Ferroportin-1 mRNA expression were measured after normalizing to β-actin expression. **E-F.** THP-1 macrophages were treated with LPS with or without 3 µM HNE or 2 µM ML385 for 3 hr followed by 6 µM nigericin (Nig) to induce inflammasome activation. (E) Cell morphology, scale bar=5 µm . (F) Cell death by SYTOX^TM^ green. Statistics in B, C, and D were performed using a 2-way ANOVA and Bonferroni’s post hoc test. **P<0.01****P<0.001 between HNE and ethanol groups after LPS treatment. Bars represent mean ± SD.

### HNE inhibits inflammasome activation independently of NF-κB signaling

Activation of the inflammasome involves first the NF-κB dependent stimulation of NLRP3 expression, and then NLRP3 oligomerization promoted by a cell danger signal, such as K^+^ efflux or ATP (36). We first focused on the role of HNE in NF-κB signaling by assessing p65 phosphorylation, p65 nuclear translocation, and IκB-α degradation, as well as the expression of TNF-α and NLRP3 in macrophages treated with LPS. In response to LPS (100 ng/mL), there was a 3-fold increase in nuclear translocation of p65 (Fig. 3A-B) and a 10-fold increase in phosphorylation of p65 (Fig. 3C). TNF-α and NLRP3 expression were induced after LPS treatment through the NF-κB pathway, but these measurements were not significantly affected by 3 µM HNE (Fig.3A-C, Fig. S2A-B). Similarly, there was no significant effect of HNE (0.3-3 µM) on NLRP3, pro-IL-1β, and IκB-α protein expression (Fig. 3D-F, and Fig. S2D ). Interestingly, HNE significantly reduced pro-IL-1β gene expression (Fig. S2C). Inflammasome activation induced by nigericin triggers IL-1β cleavage in LPS-primed macrophages. To minimize the effects of HNE on NF-κB-dependent transcription, macrophages were treated with LPS for 3 hr before exposure to HNE (Fig. S2E) and then stimulated with nigericin (Fig. 3G). We found HNE still significantly prevented IL-1β maturation (Fig. 3H). Furthermore, we performed the experiments where BMDMs were pre-treated with HNE for 30 min, and then stimulated with nigericin after 10 min LPS priming. NLRP3 activation was confirmed by the cleavage of caspase-1 and GSDMD. We found that rapid NLRP3 priming followed by nigericin treatment induced caspase-1 and GSDMD cleavage, and HNE pre-treatment inhibited these effects (Fig. S3). Thus, the inhibitory effect of HNE treatment on inflammasome activation is independent of the NF-κB pathway.

**Fig. 3.**
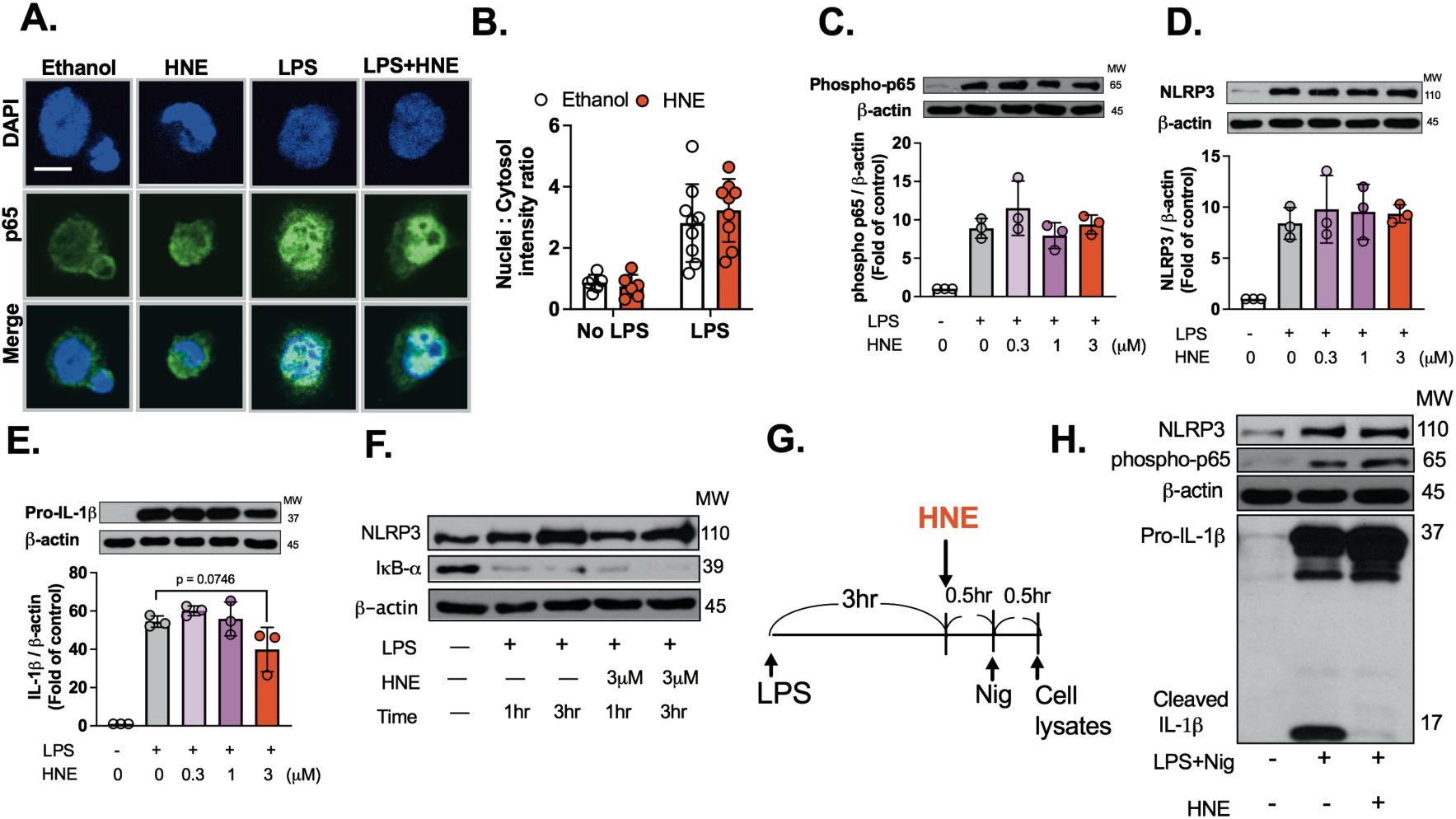
HNE inhibits IL-1β cleavage independently of NF-κB signaling in macrophages. **A-B**. THP-1 macrophages were stimulated with or without LPS (100 ng/mL) and co-incubated with ethanol or 3 µM HNE for 1 hr. (A) NF-κB p65 translocation was measured by immunofluorescence (scale bar: 5 µm), DAPI (blue), and p65 (Green). (B) Quantification using Image J. N=3 experiments **C-E.** Peritoneal macrophages were stimulated with or without LPS (100 ng/mL) and co-incubated with ethanol or HNE (0.3-3 µM) for 3 hr. Protein expression was analyzed by western blot. (C) phosphorylation of p65, (D) NLRP3, (E) Pro-IL-1β. Bars represent mean±SEM. N=3 experiments. **F.** Peritoneal macrophages were stimulated with LPS (100 ng/mL) and co-incubated with ethanol or HNE (3 µM) for 1or 3 hr. Cell lysates were analyzed by western blot. (F) NLRP3, IκB-α, and β-actin western blots are representative of three independent experiments. **G-H.** (G) Schematic of experimental design for data in Fig. 3H. (H) BMDMs were stimulated with LPS (100 ng/mL) for 3 hr followed by 6 µM nigericin (Nig) for 30 min. 3 µM HNE or ethanol was added 30 min before nigericin. Western blots (NLRP3, p65 phosphorylation, β-actin, pro-IL-1β, and cleaved IL-1β) are representative of three independent experiments. Statistics in B were performed using a 2-way ANOVA and Bonferroni’s post hoc test. Statistics in C-E were performed using a one-way ANOVA and Bonferroni’s post hoc test. N=3 experiments. A p-value less than 0.05 is statistically significant among treatment groups. Bars represent mean ± SD.

### HNE inhibits NLRP3 inflammasome activation, but has no effect on AIM2 or NLRC4

We next studied the effect of HNE on inflammasome assembly and activation. It has been suggested that NLRP3 and apoptosis-associated speck-like (ASC) proteins are assembled by acetylated α-tubulin-mediated transport to the microtubule organizing center (MTOC) (37). Upon inflammasome activation by nigericin stimulated potassium efflux, we found increased α-tubulin acetylation in LPS primed peritoneal macrophages (Fig 4A), BMDMs (Fig. S4A), and THP1 macrophages (Fig. S4B). However, HNE had no effect on acetylation of α-tubulin in these cells (Fig. 4A and Fig. S4A-B). These data suggest that acetylated α-tubulin was not involved in HNE inhibition of pyroptosis.

**Fig. 4.**
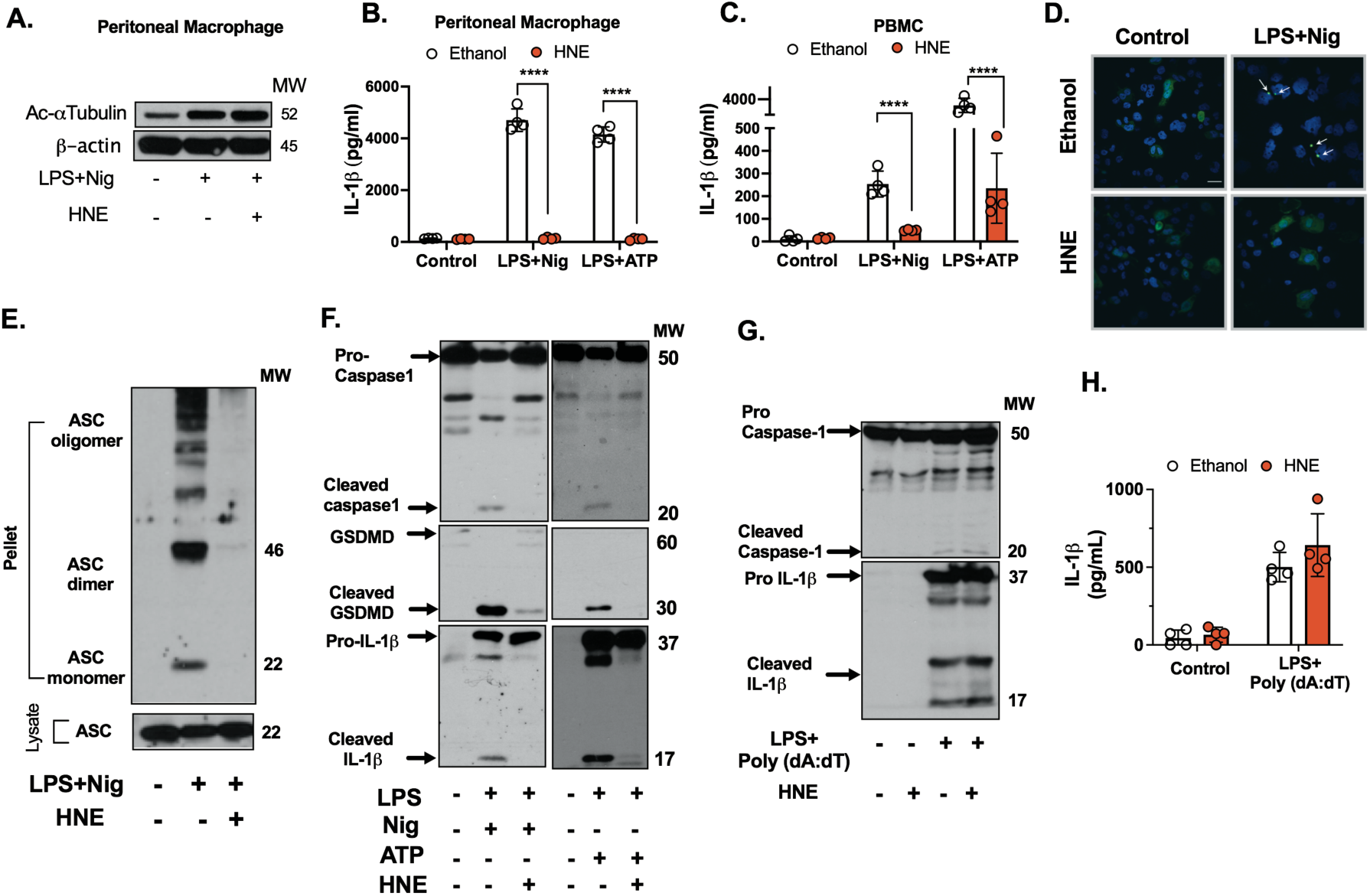
HNE inhibits NLRP3 inflammasome activation. **A-F**. HNE (3 µM) or ethanol was added 30 min before nigericin, ATP, or poly(dA:dT) treatment. **A.** Peritoneal macrophages were stimulated with LPS (100 ng/mL) for 3 hr followed by 2 µM nigericin (Nig). Acetyl-α-Tubulin (Lys40) western blots are representative of three independent experiments. **B-C.** (B) Peritoneal macrophages and (C) Human PBMC were stimulated with LPS (100 ng/mL) for 3 hr followed by 2 µM nigericin (Nig) or ATP (2 mM) stimulation for 1 hr. IL-1β in the medium was measured by ELISA. Bars represent mean±SD. **D.** THP-1 macrophages that overexpressed ASC-GFP were stimulated with LPS (100 ng/mL) followed by 6 µM nigericin (Nig) for 2 hr. ASC speck formation (arrows) was measured by confocal microscopy. (scale bar: 10 µm ). Quantification results were shown at lower magnification images in Fig. S6. **E.** THP-1 macrophages were stimulated with LPS (100 ng/mL) for 3 hr followed by 6 µM nigericin (Nig) for 2 hr. ASC oligomerization western blots are representative of three independent experiments. **F.** Peritoneal macrophages were stimulated with LPS (100 ng/mL) for 3 hr followed by 2 µM nigericin (Nig) or ATP (2 mM) stimulation for 15 min. Western blots are representative of three independent experiments. **G.** Peritoneal macrophages were stimulated with LPS (100 ng/mL) for 3 hr followed by 2µg/mL Poly(dA:dT) for 6 hr. Ethanol or 3 µM HNE was co-incubated with cells 30 min before Poly(dA:dT). Western blots are representative of three independent experiments. **H.** THP-1 macrophages were stimulated with LPS (100 ng/mL) for 3 hr followed by 2µg/mL Poly(dA:dT) for 6 hr. IL-1β in the medium was measured by ELISA. Statistics in B, C, and H were performed using a 2-way ANOVA and Bonferroni’s post hoc test. ****P<0.001 between control and treatment groups. Bars represent mean ± SD.

To analyze the effects of HNE more broadly, we used both ATP and nigericin as a second signal to induce inflammasome activation (Fig. 4B-C). We also studied both peripheral blood mononuclear cells (PBMC) from healthy human donors, and peritoneal macrophages from mice. We confirmed that HNE inhibited IL-1β release (Fig. 4B-C).

Next, the effect of HNE on a non-potassium efflux dependent NLRP3 stimulus was tested. THP-1 macrophages were stimulated with LPS for 3 hr followed by R837 (Imiquimod) for 1 hr. We found that HNE reduced IL-1β secretion (Fig. S5). These data indicated that HNE also inhibits non-potassium efflux-mediated NLRP3 activation.

It is known that upon NLRP3 inflammasome activation, the adaptor protein ASC is recruited by NLRP3 and forms large multimeric complexes, termed ASC specks (38). To determine the effect of HNE on ASC speck formation, we overexpressed an ASC-GFP fusion protein in THP-1-differentiated macrophages and stimulated them with LPS followed by nigericin. HNE treatment inhibited ASC speck formation, as indicated by reduced ASC speck immunofluorescence (Fig. 4D, and Fig. S6A-B). Similarly, HNE reduced large multimeric ASC complexes, as detected by western blot (Fig. 4E).

Inflammasome activation leads to the cleavage of pro-caspase-1 to generate active caspase-1 that cleaves gasdermin-D (GSDMD) to form membrane pores, which enables the release of cytokines, such as IL-1β. Therefore, we measured the effect of HNE on these parameters of inflammasome activation (Fig. 4F). LPS primed peritoneal macrophages were treated with HNE followed by nigericin or ATP. The inhibitory effect of HNE on NLRP3 activation was confirmed by western blot as shown by decreased cleavage of caspase-1, GSDMD, and IL-1β to their mature p20, p30, and p17 forms, respectively (Fig. 4F).

NLRP3, AIM2, and NLRC4 inflammasomes respond to different ligands or activators, but all engage with the adaptor protein ASC and activate protein caspase-1 to cleave pro-IL-1β. To study the specificity of HNE for the NLRP3 inflammasome, we tested the effect of HNE on the AIM2 and NLRC4 inflammasome. AIM2 is activated by cytosolic double-stranded DNA (dsDNA). NLRC4 is activated by cytosolic flagellin. To determine the effect of HNE on these two inflammasomes, macrophages were treated with LPS for 3 hr, then HNE for 30 minutes, then transfection with poly(dA:dT) or flagellin for 6 hr and 3 hr, respectively. In contrast to the NLRP3 inflammasome, HNE did not reduce cleavage of caspase-1 or IL-1β release after AIM2 inflammasome activation (Fig. 4G-H). Similarly, HNE did not reduce IL-1β release after NLRC4 inflammasome activation (Fig. S7). Together, these data show that HNE inhibits NLRP3 assembly and activation in mouse and human cells, but has no effect on AIM2 and NLRC4.

### HNE reduces non-canonical inflammasome-mediated IL-1β release, but has no effect on GSDMD-mediated cell death

Non-canonical inflammasome activation is dependent on caspase-11 (in mice) or caspase-4 (in humans) which directly bind with intracellular LPS (39, 40). Recent data suggest that the non-canonical inflammasome pathways play critical roles in acute lung injury and sepsis by sensing cytosolic LPS (41). To test whether HNE also affects non-canonical inflammasome activation, we transfected LPS into peritoneal macrophages in the presence or absence of HNE. IL-1β released into the medium and within whole cell lysates was collected 16 hr after transfection. Release of IL-1β into the medium and cleavage of IL-1β in the cell lysates were decreased 50% by HNE compared to control, but NLRP3 and pro-IL-1β protein levels did not change (Fig. 5A-B). Furthermore, HNE had no effect on Gasdermin-D (GSDMD) cleavage, but partially reduced IL-1β maturation (Fig. 5B). Although caspase-11 is the major enzyme that cleaves GSDMD during non-canonical inflammasome activation, IL-1β maturation remains dependent on canonical NLRP3 activation (11). These data suggest that HNE targets NLRP3, but not caspase-11.

**Fig. 5.**
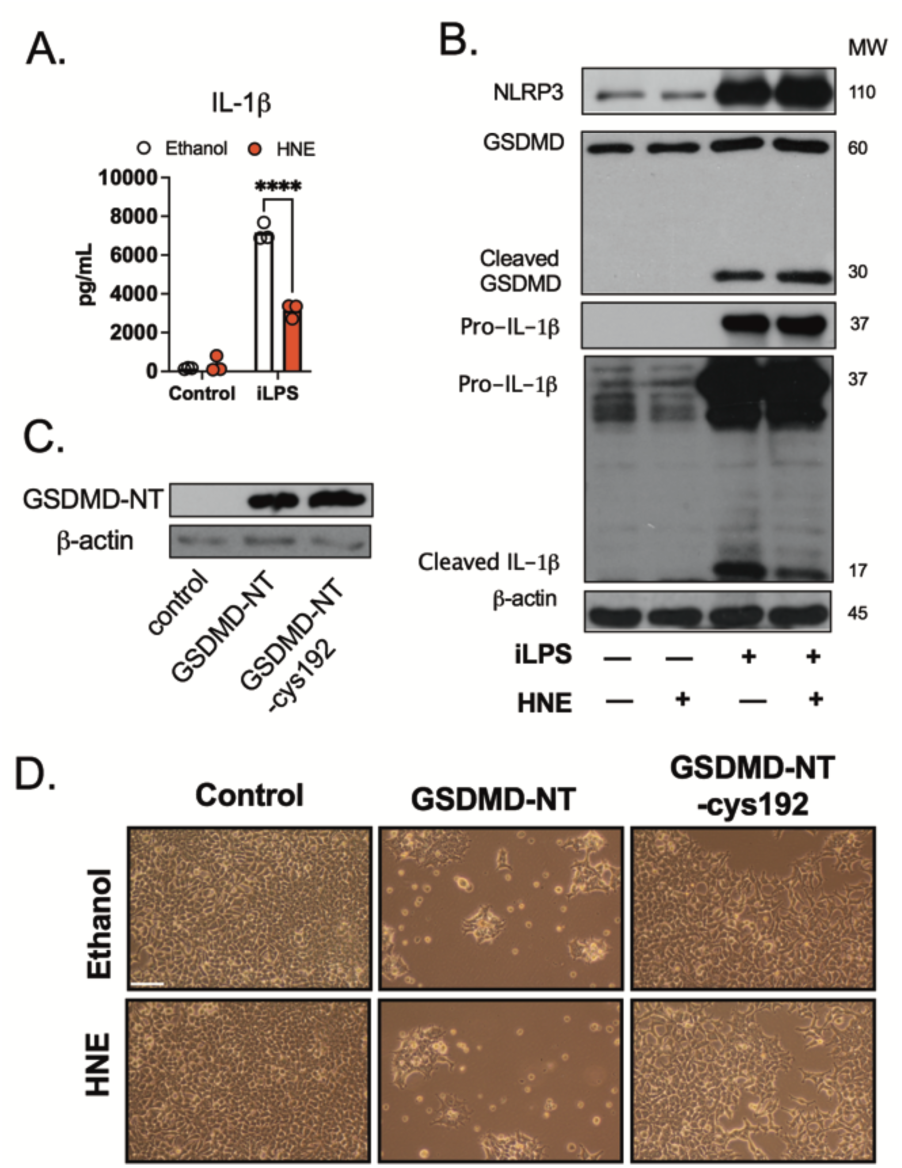
HNE inhibits non-canonical inflammasome-mediated IL-1β secretion, but has no effect on GSDMD-mediated cell death. **A-B**. Peritoneal macrophages were stimulated with LPS (100 ng/mL) for 3 hr followed by intracellular LPS transfection (1µg/mL) for 16 hr. Ethanol or 3 µM HNE was co-incubated with cells 30 min before intracellular LPS transfection. (A) IL-1β in the medium (B) Western blots are representative of three independent experiments. **C-D.** HEK293 cells were transfected with GSDMD-NT or GSDMD-NT-cys192 to Ala192. Ethanol or 3 µM HNE was co-incubated with cells. (C) Western blots are representative of three independent experiments. (D) Micrographs of cultures were obtained 48 hr after transfection under phase contrast illumination using 20X objective, scale bar=100 µm. Images are representative of three independent experiments. Statistics in A were performed using a 2-way ANOVA and Bonferroni’s post hoc test. ****P<0.001 between control and treatment groups. Bars represent mean ± SD.

The assembly of the GSDMD pore requires cleavage of an autoinhibitory sequence present in the GSDMD-C-terminal domain (GSMDM-CT). This allows the N-terminal domain (GSDMD-NT) to bind to the inner leaflet of the plasma membrane, where it oligomerizes, and forms pores. Recent data show that Cys192 (Cys191 in humans) is critical for GSDMD-NT oligomerization (42). To test the possibility of a HNE effect on GSDMD-NT, and subsequent GSDMD-mediated cell death, we overexpressed GSDMD-NT in HEK293T cells. HNE was added after transfection. Overexpression of GSDMD-NT stimulated cell death, but HNE treatment did not protect the cells (Fig. 5C-D). Consistent with previous studies (42, 43), GSDMD-NT-C192A reduced cell death (Fig. 5D). Together, these results suggest that HNE specifically inhibits the NLRP3 inflammasome, but not GSDMD cleavage associated cell death.

### HNE inhibits inflammasome activation by blocking the NLRP3-NEK7 interaction via a cysteine-dependent mechanism

The family of mammalian NIMA-related kinases 7 (NEK7) was recently identified as an NLRP3-binding protein that regulates NLRP3 oligomerization and activation (44-47). To detect a physical association between NEK7 and NLRP3, we performed co-immunoprecipitation assays. We found NEK7 was associated with NLRP3 after LPS/nigericin treatment, but the association was inhibited by HNE treatment (Fig, 6A and Fig. S8A-D). These results demonstrated that HNE reduced NLRP3 inflammasome activation by blocking the interaction between NLRP3 and NEK7.

To determine if HNE binds directly to NLRP3, we used a click chemistry-based approach. For this, we treated cells with alkyne-HNE (48), which can be activated by azido click chemistry to bind biotin. First, we compared the effects of HNE and alkyne-HNE on inflammasome activation. Like HNE, alkyne-HNE inhibited IL-1β release upon NLRP3 inflammasome activation in a dose-dependent manner in peritoneal macrophages (Fig. 6B-C). Alkyne-HNE was 3 times less potent than HNE as measured by IL-1β release (3 µM HNE vs 10 µM alkyne-HNE). As anticipated, when LPS-primed macrophages were treated with alkyne-HNE, the cleavage of GSDMD and IL-1β were both inhibited in a dose dependent manner, as shown by increased pro-IL-1β, decreased GSDMD and cleaved IL-1β (Fig. 6D). We next assessed if alkyne-HNE can bind to NLRP3 by treating THP-1 macrophages with 3 µM HNE or 10 µM alkyne-HNE after inflammasome activation by nigericin. There were no significant differences in NLRP3 protein expression from whole cell lysates between HNE or alkyne-HNE groups.

**Fig. 6.**
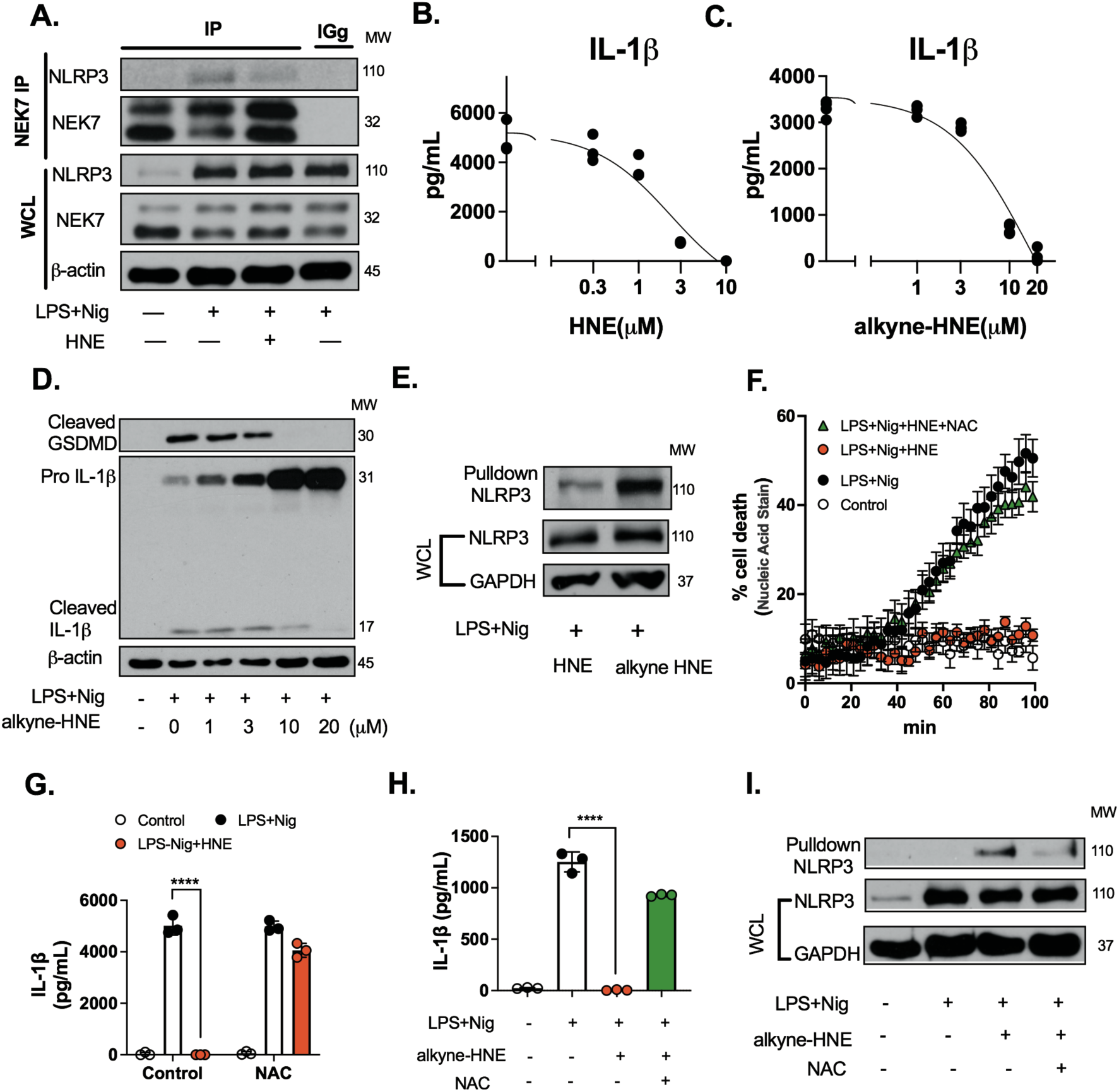
HNE inhibits inflammasome activation by blocking the NLRP3-NEK7 interaction and a cysteine dependent mechanism. **A** BMDM were stimulated with LPS (100 ng/mL) for 3 hr followed by 2 µM nigericin (Nig) for 60 min. Ethanol or 3 µM HNE was added 30 min before nigericin. Immunoblots of NLRP3, NEK7, and β-actin from NEK7-immunoprecipitated and whole cell lysates (WCL) are representative of three independent experiments. **B-D.** Peritoneal macrophages were stimulated with LPS (100 ng/mL) for 3 hr followed by 2 µM nigericin (Nig) for 1 hr. HNE (0-10 µM) or alkyne-HNE (0-20 µM) was added 30 min before nigericin. (B and C) IL-1β in the medium. The dose-response curve was plotted by using a logarithmic x-axis to cover a range of HNE or alkyne-HNE concentrations. (D) Protein expression was measured by western blot. (N=3-4 independent experiments) **E.** THP-1 macrophages were stimulated with LPS (100 ng/mL) for 3 hr followed by 6 µM nigericin (Nig) for 30 min. HNE (3 µM) or alkyne-HNE (10 µM) was added 30 min before nigericin. Immunoblots of NLRP3 pulldown with streptavidin after performing a click reaction on whole cell lysates (WCL). Data are representative of three independent experiments. **F-G.** THP-1 macrophages were stimulated with LPS (100 ng/mL) for 3 hr followed by nigericin (Nig, 6 µM) for 2 hr. HNE (3 µM), n-acetyl cysteine (NAC, 500 µM), or both were co-incubated with cells. (F) Cell death by SYTOX^TM^ green. (G) IL-1β in the medium. **H-I.** BMDM were stimulated with LPS (100 ng/mL) for 3 hr followed by nigericin (Nig, 2 µM) for 30 min. Alkyne-HNE (10 µM), N-acetyl cysteine (NAC, 500 µM), or both were added 30 min before nigericin. (H) IL-1β in the medium. (I) Immunoblots of NLRP3 pulldown with streptavidin after performing a click reaction on whole cell lysates (WCL) are representative of three independent experiments. Statistics in G and H were performed using a 2-way ANOVA and Bonferroni’s post hoc test. ****P<0.001 between control and treatment groups. Bars represent mean ± SD.

However, by using alkyne-HNE and the azido click-chemistry technique, followed by a biotin-streptavidin pulldown, we found that alkyne-HNE pulled down significantly more NLRP3 than HNE treatment (Fig. 6E and Fig. S9). These data suggest that alkyne-HNE was capable of targeting NLRP3 during inflammasome activation.

HNE is highly electrophilic and may interact with NLRP3 cysteines by covalent modification. Indeed, when HNE or alkyne-HNE was co-incubated with N-acetylcysteine (NAC) that contains a reactive cysteine, which can inactivate cysteine-reactive metabolites, the ability of HNE to protect cells from nigericin-mediated cell death, IL-1β release and MitoSOX nuclear accumulation was eliminated (Fig. 6F-H, and Fig. S10A-B). Furthermore, western blots indicated that alkyne-HNE-NLRP3 interaction was significantly decreased in the presence of NAC (Fig. 6I). 4-hydroxynonenal glutathione (HNE-GSH) is a major product formed by the reaction of HNE with GSH. To test if the reduced HNE is able to inhibit pyroptosis, we used LPS co-incubated with HNE-GSH, followed by nigericin in THP-1 macrophages. Cell death was confirmed by morphological changes, and LDH release. We found that HNE-GSH treatment alone (3 and 10 µM) had no effect on cell death compared to LPS/nigericin-treated cells (Fig. S11). These findings suggest that the interaction between HNE and NLRP3 is mediated by a cysteine dependent mechanism.

### HNE inhibits inflammasome activation in acute lung injury (ALI) and sepsis models

To test the ability of exogenous HNE to inhibit lung injury and inflammasome activation in vivo, we used the LPS induced ALI model (49-51). C57BL/6 mice were exposed to a single dose of saline, HNE, LPS or LPS+HNE by oropharyngeal delivery. Lung tissue and bronchoalveolar lavage (BAL) fluid were collected 3 hr and 18 hr after exposure to assess lung injury and the inflammatory response using myeloperoxidase (MPO) staining, Ly6G&6C staining, serum amyloid A-3 (SAA3) mRNA expression, cleaved IL-1β immunohistochemistry (IHC), as well as IL-1β and TNF-α secretion in BAL fluid. LPS delivery caused intense neutrophil and macrophage infiltration in the lung, and increased IL-1β staining (Fig. 7A-D, Fig. S12A). LPS delivery also stimulated IL-1β secretion in the BAL fluid and total IL-1β production in lung homogenates (Fig. 7E-F). Simultaneous delivery of HNE with LPS significantly decreased all these markers of inflammasome activation compared to LPS alone (Fig. 7B-G), but had no effect on TNF-α (Fig. S12B) supporting that HNE was specific for NLRP3 inflammasome.

**Fig. 7.**
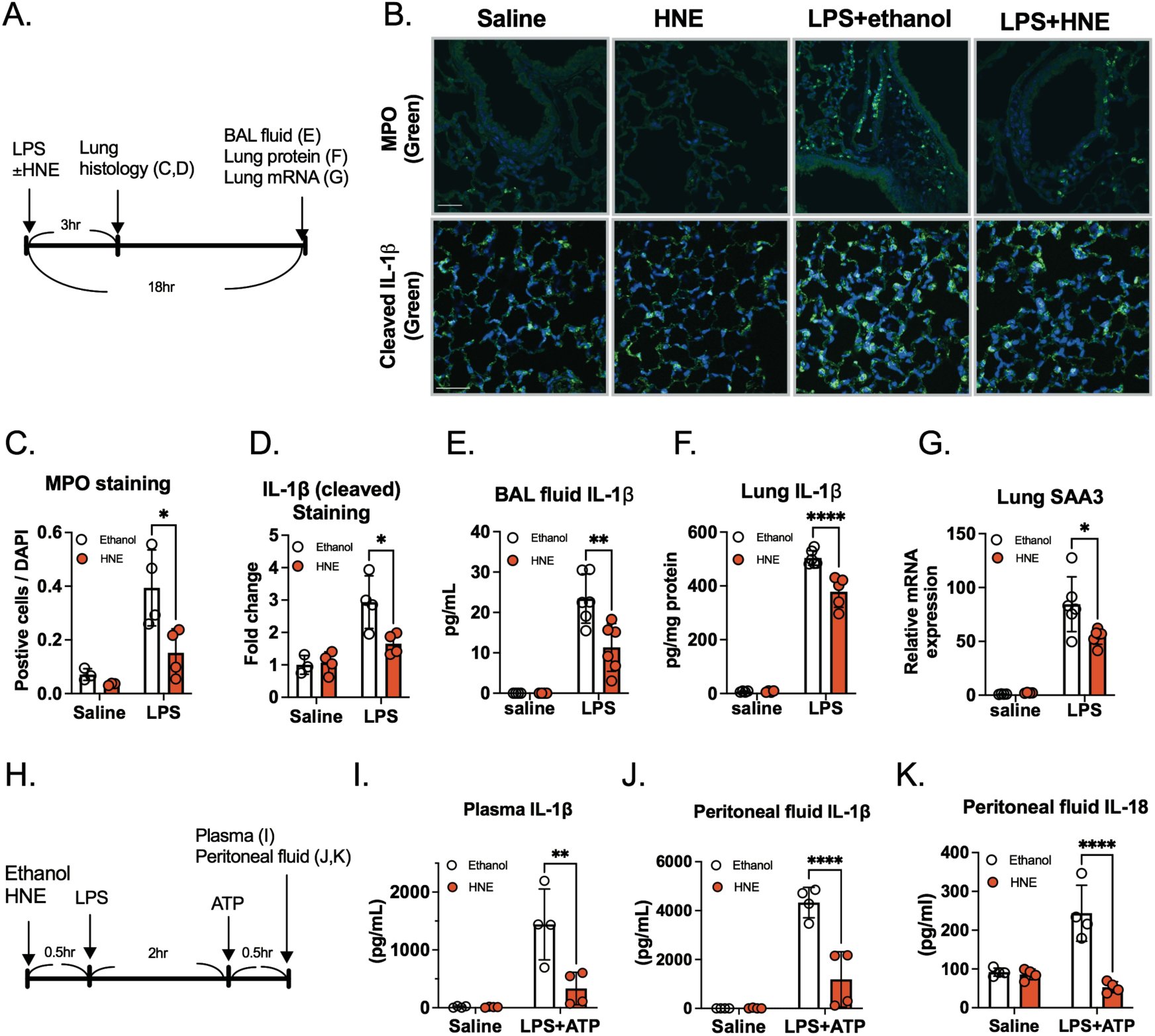
HNE inhibits IL-1β and IL-18 secretion in mouse acute lung injury model and sepsis model. **A-G**. (A) Schematic of experimental design for LPS-induced acute lung injury in mice. Ethanol (0.01%) in saline (50 µl), HNE (6 µM) in saline (50 µl), LPS (2 mg/kg) + ethanol (0.01%) in saline, or LPS (2 mg/kg) + HNE (6 µM) in saline were delivered oropharyngeally to mice. Lung tissues were harvested at 3 hr post treatment for immunohistochemistry (N=4 mice from each group). BAL fluid, lung tissue protein and RNA were harvested at 18 hr (N=4 mice from each saline group, and N=6 from each LPS group). (B) Immunofluorescence of representative lung sections for myeloperoxidase (MPO) (Green) or cleaved IL-1β (Green). DAPI (Blue), (scale bar=40 µm). Quantification results of (C) MPO, and (D) cleaved IL-1β. (E) IL-1β protein in BAL fluid and (F) IL-1β protein in lung tissue. (G) SAA3 mRNA expression in lung tissue was measured by real-time PCR and normalized to β-actin. **H-K.** (H) Schematic of experimental design for LPS-ATP-induced inflammasome activation in mice. Mice were pre-injected with ethanol, HNE (2 mg/kg) 0.5 hr before LPS (10 mg/kg) i.p. for 2 hr followed by ATP (100 mM in 100 µl, pH 7.4) i.p. Plasma and peritoneal fluid were harvested 0.5 hr after ATP treatment for cytokine assay measured by ELISA . (I) IL-1β in plasma, (J) IL-1β in peritoneal fluid. (K) IL-18 in peritoneal fluid. (N=4 mice from each group). Statistics in C-G and I-K were performed using a 2-way ANOVA and Bonferroni’s post hoc test. *P<0.05, **P<0.01, ****P<0.001 between ethanol and HNE treatment groups after injury. Bars represent mean ± SD.

Sepsis is characterized by multi-organ dysfunction caused by an exaggerated immune response to infection (52). We characterized a clinically relevant, acute in vivo sepsis model to study the inflammasome pathway by using a combination of LPS and ATP (53). To test whether exogenous HNE could inhibit inflammasome activation in this model, C57BL/6 mice were injected with vehicle or HNE, 30 min before receiving LPS (10mg/kg, i.p.). Two hours later, mice were injected with ATP (Fig. 7H). Plasma and peritoneal fluid were harvested 30 min after ATP injection for measurement of IL-1β and IL-18. Administration of HNE significantly reduced the levels of both cytokines (Fig. 7I-K). Collectively, these data show that HNE treatment reduces inflammasome activation in vivo in both ALI and sepsis models.

### Increasing endogenous HNE by the GPX4 inhibitor RSL3, reduces inflammasome activation in macrophages in vitro and in vivo

HNE is one of the most abundant lipid peroxidation products (54). Excessive ROS reacts with the polyunsaturated fatty acids of lipid membranes and induces lipid peroxidation. Glutathione peroxidases (GPXs) are antioxidant enzymes that protect cells from lipid peroxidation (55). It is established that glutathione peroxidase-4 (GPX4) deficiency or inhibition enhances the production of endogenous HNE in vivo and in vitro (56-58). Indeed, the GPX4 inhibitor, RSL3, induced a dose-dependent increase of HNE in BMDMs (Fig. S13). To assess the effect of endogenous HNE in pyroptosis, we treated THP-1 macrophages and peritoneal macrophages with RSL3, after LPS-nigericin stimulation. RSL3 treatment significantly reduced cell death (Fig. 8A-B). RSL3 also inhibited GSDMD cleavage and IL-1β secretion in a dose-dependent manner (Fig. 8C-D). To test whether RSL3 treatment also inhibited inflammasome activation in vivo, C57BL/6 mice were injected with vehicle, or RSL3 (2mg/kg) before LPS and ATP challenge. Plasma and peritoneal fluid were harvested 30 min after ATP for measurement of IL-1β. We found that administration of RSL3 significantly reduced the levels of IL-1β in plasma and peritoneal fluid (Fig. 8E-F). Collectively, these data indicate that the accumulation of HNE, induced by GPX4 inhibition, decreased inflammasome activation in vitro macrophages and in a mouse sepsis model.

**Fig. 8.**
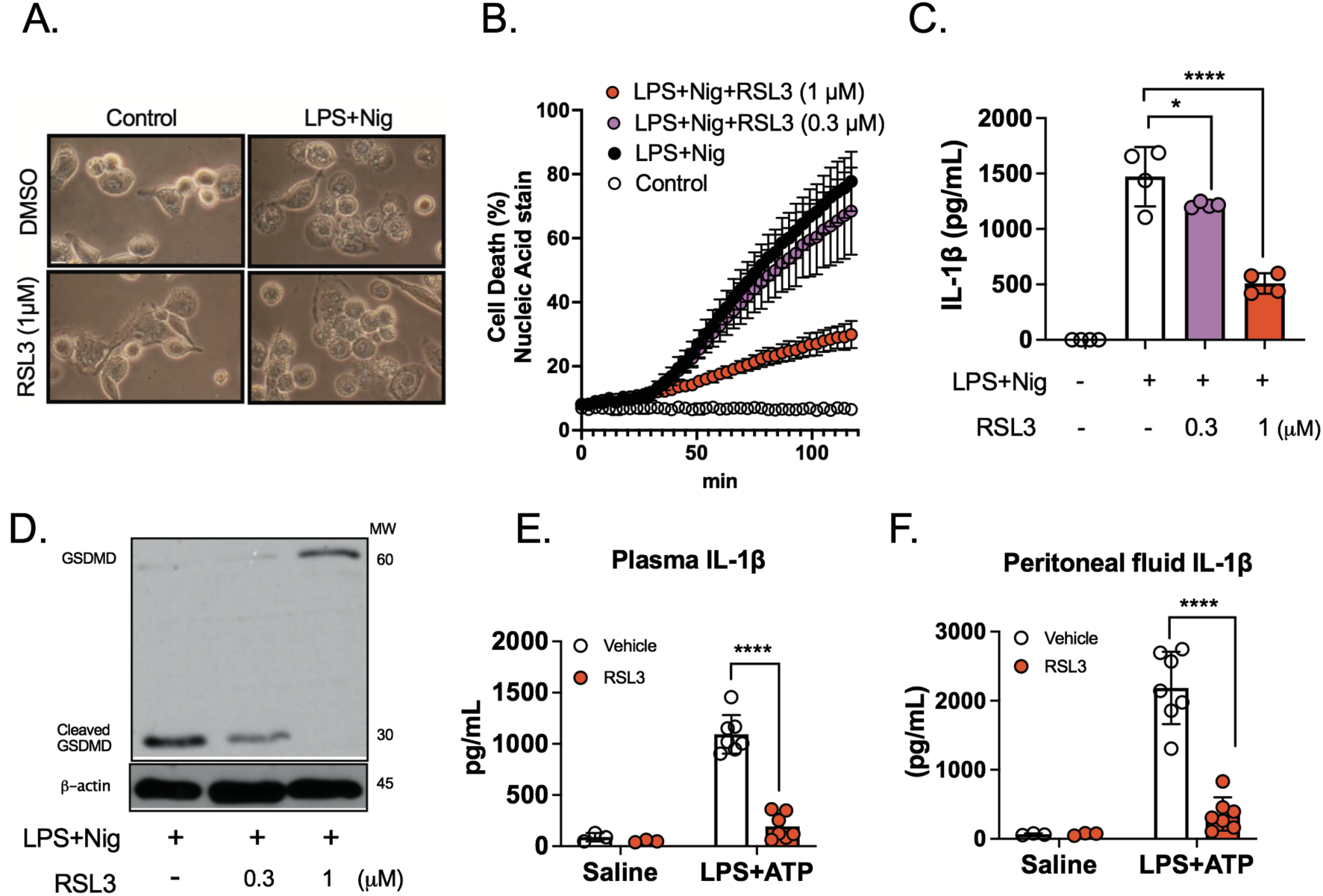
GPX4 inhibition by RSL3 reduces inflammasome activation in macrophages and in an in vivo sepsis model. **A-B**. THP-1 macrophages were stimulated with LPS (100 ng/mL) for 3 hr followed by nigericin (Nig, 6 µM) for 2hr. DMSO, or RSL3 (0.3 or 1 µM) was co-incubated with cells. (A) Cell morphology, scale bar= 5 µm . (B) Cell death by SYTOX^TM^ green. **C-D.** Peritoneal macrophages were stimulated with LPS (100 ng/mL) with or without RSL3 (0.3 or 1 µM) for 3 hr followed by nigericin (Nig, 2 µM) for 30 min. (C) IL-1β in the medium. (D) Western blots are representative of three independent experiments. **E-F.** Mice were injected i.p. with vehicle, or 2 mg/kg RSL3 30 min before LPS (10mg/kg). Two hr after LPS, mice were injected i.p. with ATP (1mg/kg). After 30 min, plasma and peritoneal fluid were harvested for ELISA. (E) plasma IL-1β, and in (F) peritoneal fluid IL-1β. N=3 mice from each saline group, and N=7 from each LPS-ATP group Statistics in C, E, and F were performed using a 2-way ANOVA and Bonferroni’s post hoc test. *P<0.05, ****P<0.001 among treatment groups. Bars represent mean ± SD.

## Discussion

The major finding in the present study was that the abundant lipid peroxidation product HNE selectively inhibited NLRP3 inflammasome activation in mouse and human macrophages. Three key results were: 1) HNE blocked NLRP3 inflammasome-mediated pyroptosis and IL-1β release in mouse macrophages and human PBMC, independent of Nrf2 and NF-κB signaling pathways. 2 HNE directly bound to NLRP3 and inhibited its interaction with NEK7. 3) HNE inhibited inflammasome activation in mouse acute lung injury and sepsis models. Our data suggest that HNE is not only a product of lipid peroxidation, but also plays an important role as a signaling molecule that inhibits inflammation at physiological concentrations (3-10 µM) by regulating NLRP3 inflammasome activation.

The mechanism we propose for the protective effect of HNE against pyroptosis is that it directly inhibits NLRP3 inflammasome activation by blocking NEK7-NLRP3 interaction. Our data in support of this mechanism include: 1) HNE treatment of LPS-primed macrophages immediately before activation of NLRP3 with nigericin or ATP reduced IL-1β release, suggesting that HNE exerts its effect after the priming step. 2) HNE blocked ASC oligomerization and speck formation which is upstream of caspase-1, GSDMD, and IL-1β cleavage. 3) HNE had no effect on IL-1β secretion mediated by AIM2 and NLRC4 inflammasome activation, although NLRP3, NLRC4, and AIM2 inflammasomes share the adaptor protein ASC and effector protein caspase-1. 4) Alkyne-HNE directly bound to NLRP3 in human and mouse macrophages. 5) HNE blocked NLRP3-NEK7 interaction. Therefore, we believe that the binding of HNE to NLRP3 is necessary and sufficient to inhibit NLRP3 inflammasome activation.

The NEK7-NLRP3 interaction is essential for NLRP3 assembly (44, 47), and subsequent ASC speck formation, caspase-1 activation, IL-1β release, and pyroptosis. As previously reported, NEK7 depletion does not affect IL-1β release in macrophages when treated with poly(dA:dT) (44, 47). These data show that NEK7 is required for NLRP3 inflammasome activation, but not AIM2 inflammasome activation (44, 47), supporting our findings that HNE specifically targets NLRP3. A recent cryo-electron microscopy study revealed that NLRP3 interacts with NEK7 at leucine-rich repeat (LRR) and NACHT domains (46). Additional research will be needed to evaluate the structural mechanism for HNE-NLRP3 interaction.

Pyroptosis is characterized by cell swelling, membrane rupture, and release of inflammatory cytokines, such as IL-1β and IL-18 (59). In contrast to our study, Kauppinen et al. and Jin et al., showed that high concentrations of HNE (30-200 µM) increased NLRP3 mRNA expression, IL-1β, and IL-18 secretion in human ARPE-19 cells after LPS stimulation (60, 61). We believe that the results obtained in these studies were due to HNE toxicity at concentrations 30-60-fold greater than in this present study. Therefore, we believe that physiological concentrations of HNE protect against pyroptosis and IL-1β release.

It has been hypothesized that ROS generation triggers NLRP3 inflammasome activation (62, 63) and pyroptosis (64). Nrf2 induction by HNE may stimulate the expression of potent antioxidant and cytoprotective proteins that increase the resistance to cytotoxic ROS. This mechanism was supported by evidence that electrophiles (such as sulforaphane, tert-butyl hydroquinone, dimethyl fumarate and itaconate) showed anti-inflammatory responses through Nrf2 pathways (65-69). However, we found that HNE inhibited pyroptosis independent of Nrf2 (Fig. 2E-F), suggesting the existence of another HNE mediated pathway for inhibition of NLRP3 inflammasome activation.

To determine the effect of HNE on cysteine reactive pathways, we measured the effects of HNE vs NAC on pyroptosis and IL-1β secretion (Fig. 6F-H), after nigericin stimulation of LPS-primed THP-1 macrophages and BMDMs. Previous studies showed that NAC (>5mM ) inhibited NLRP3 responses in macrophages (70, 71), contradicting our data (Fig. 6F-H). We think the differences between these studies and our study is that we used only 0.5 mM NAC which had no effect on pyroptosis and IL-1β secretion. It has been shown that high concentration of NAC (> 5mM) inhibited transcription of NLRP3, but had no effect on activation (72, 73). Our results are supported by a recent publication indicating that 0.5 mM NAC alone had no effect on cell death after LPS+nigericin treatment, but was sufficient to reverse the protective effect of another cysteine-reactive drug, disulfiram, on pyroptosis (43). In our study, we found 0.5 mM NAC was sufficient to reverse the HNE effect suggesting that HNE inhibits NLRP3 inflammasome through a cysteine dependent mechanism.

Previous studies suggested a role for HNE in regulating IL-1β secretion via NF-κB signaling in monocytes and macrophages when cells were treated with high dose HNE (25 µM) prior to LPS stimulation (26, 74). In our study, neither translocation or phosphorylation of p65, nor protein expression of NLRP3 and IκB-α were altered by HNE treatment (0.3-3 µM) after LPS stimulation (Fig. 3), suggesting a limited role for NF-κB. It is worth noting in previous studies that IL-1β secretion was extremely low, and there was no pyroptosis, because a second signal to induce NLRP3 activation was not included (26, 60, 74); which we have shown using nigericin or ATP.

Although transcriptional priming is required for NLRP3-mediated IL-1β secretion, our data (Fig. 3, and Fig. S3) and other studies have demonstrated that transcription is not necessary for NLRP3 activation (75, 76). Instead, posttranslational modifications including ubiquitination, phosphorylation, dephosphorylation and many other processes are essential for NLRP3 activation (77). For example, cyclic AMP promotes NLRP3 ubiquitination through the E3 ubiquitin ligase MARCH7 (78), and protein kinase A (PKA) directly phosphorylates Ser295 in NLRP3, which is critical for NLRP3 oligomerization (79). It is unclear how HNE affects the posttranslational modifications of NLRP3. Future studies on the mass spectrometric analysis of the NLRP3 inflammasome should identify the detailed mechanisms.

NLRP3 inflammasome activation plays an important role in the pathogenesis of acute lung injury (ALI) and sepsis (50, 52, 80). In the present study, we found that co-delivery of HNE with LPS to the lungs significantly reduced IL-1β cleavage and inflammatory cell infiltration. An important aspect of our work is that HNE is primarily an endogenous product of lipid peroxidation (16, 17). To determine the endogenous effect of HNE, we increased its levels in cells by inhibiting GPX4 (56-58). We found that peritoneal administration of HNE or the GPX4 inhibitor, RSL3, decreased IL-1β levels in peritoneal fluid and plasma after LPS-ATP challenge. These results suggest that both exogenous and endogenous HNE can inhibit NLRP3 inflammasome activation. A previous study by Kang et al. found that depletion of GPX4 fro myeloid cells increased septic lethality (81), which contradicts our findings. It is possible that chronic GPX4 deficiency may cause cell death due to accumulation of peroxidation, increased mitochondrial DNA and cytosolic ATP release, thereby triggering inflammasome activation and cell death.

In summary, our data indicate that HNE is not just a pathogenic mediator of oxidative stress (18, 54), but a novel endogenous inhibitor of NLRP3 inflammasome activation and subsequent inflammation. Regulation of HNE formation may represent a new therapeutic approach to inhibiting NLRP3 inflammasome activation, IL-1β secretion, and tissue inflammation.

## Materials and Methods

### Animal

Female and male C57BL/6J mice (Jackson Laboratory, 000664) aged 10-14 weeks were used in the experiments. All mice were housed in a specific pathogen-free facility and kept in a temperature-controlled room set to a light and dark cycle of 12 hours each. The mice had ad libitum access to standard mouse chow and water. No sample or animal was excluded from the experiments. Experiments with animals were blinded to the researcher assessing markers.

### Oropharyngeal administration of LPS and HNE

Animals were randomly allocated to experimental groups. Age matched mice were anesthetized with ketamine (40 mg/kg) /xylazine (3 mg/kg). After respiratory rate was significantly decreased, the mouse was suspended by a rubber band on a 60° incline board. The oral cavity was exposed and the tongue was fully extended by forceps. Using a P200 micropipette, 50 µl physiological saline or 2mg/kg lipopolysaccharides (LPS) from Escherichia coli O127:B8, LPS (Sigma-Aldrich, L3129) with ethanol or 6µM HNE (Millipore Sigma, 393204) diluted in saline was instilled into the oropharyngeal space of mice. The nose was occluded and the tongue extended for 5 seconds following clearance of the liquid from the oropharynx. Animals were subsequently removed from the board and observed closely until fully recovered from anesthesia.

### Collection of bronchoalveolar lavage (BAL) fluid and lung tissue for protein and RNA

Mice were euthanized via intraperitoneal injection of ketamine (130 mg/kg) and xylazine (8.8mg/kg) before euthanasia. The upper part of the trachea was cannulated and then lavaged with 1 mL followed by 0.5 mL of PBS supplemented with 1 mM UltraPure EDTA (Thermo Fisher Scientific, 15575-038). BAL fluid was centrifuged at 500 x g for 5 min at 4°C and the supernatants were analyzed for cytokines and chemokines. The superior lobe, and inferior lobe were snap frozen in liquid nitrogen to quantify gene and protein expression. The superior lobe was homogenized in RTL lysis buffer (QIAGEN, 74106) and the inferior lobe was homogenized in cell lysis buffer (Cell Signaling Technology) supplemented with 5 mM sodium fluoride (Sigma-Aldrich) and protease inhibitor cocktail (Sigma-Aldrich. P8340). Protein levels were measured by Pierce™ BCA Protein Assay Kit (Thermo Fisher Scientific, 23225).

### Collection of lung tissue for immunofluorescent staining

Mice were euthanized via intraperitoneal injection of ketamine (130 mg/kg) and xylazine (8.8mg/kg). The lungs were perfused free of blood by gentle infusion of 10 ml PBS containing 1 mM EDTA through the right ventricle. Lungs were inflated with 2mL, PBS-equilibrated 4% formaldehyde (VWR International, PI28908). Lung tissues were removed, drop fixed in 4% formaldehyde, embedded in paraffin, cut into 5 μm sections, and mounted onto slides. Sections were deparaffinized before use. Sections were washed 3 times in PBS followed by antigen retrieval for 20 minutes with steam using 1X Citrate buffer (Millipore, 21545), pH=6.0. Sections were blocked in 10% normal goat serum (Vector Laboratories, S-1000) in PBS for 1 hour at room temperature followed by overnight incubation at 4°C with MPO antibody (Thermo Fisher Scientific, PA5-16672) 1:500, cleaved IL-1β antibody (Thermo Fisher Scientific, PA5-105048) 1:500, or Ly6G&6C antibody (BD Biosciences, 550291) in 2% normal goat serum in PBS overnight. After three washes with PBS , fluorescence-conjugated secondary antibodies (Molecular Probes, A-11034 or A-11081 ) 1:1000 were incubated for 1 hour at room temperature and followed by three washes with PBS. Nuclei were stained with DAPI-fluoromount-G (Southern Biotech, 0100-20). Fluorescent images were captured using a confocal microscope (Olympus BX51, Software: SPOT Imaging software advanced). MPO, IL-1β, Ly6G&6C and DAPI positive cells were quantified by using NIH ImageJ software.

### In vivo sepsis model

Sepsis was induced in C57BL/6J mice by intraperitoneal (IP) injection with LPS (10 mg/kg) for 2 hr followed by 100 mM/100µl, pH 7.4 ATP (Sigma-Aldrich, A7699) IP injection as previously described (53). Animals were randomly allocated to experimental groups. Mice were IP injected with ethanol, HNE (2 mg/kg), DMSO, or RSL3 (2mg/kg) (Cayman Chemical Company, 19288) 0.5 hr before LPS. Blood samples and peritoneal fluid were collected at 30 min after ATP injection. Whole blood samples were collected in tubes with 10 µL 500 mM EDTA and centrifuged at 1000 g for 10 min at 4°C. Plasma was collected and aliquoted for cytokine assay. Peritoneal fluids were harvested by washing mouse peritoneal cavity with 5 mL ice-cold PBS supplemented with 1mM EDTA. The cell suspension was centrifuged at 500 g for 5 min at 4°C, and the supernatant was collected and aliquoted for cytokine assay.

### Peritoneal macrophage isolation

Peritoneal macrophage isolation was performed as previously described (51, 82). One ml of sterile Bio-Gel P-100 polyacrylamide beads (Bio-Rad, 150-4174) in PBS (2% w/v) was injected IP into male and female C57BL/6J. Four days after injection, the animals were euthanized by CO_2_ and macrophages were harvested by washing their peritoneal cavity with 7 mL ice-cold PBS supplemented with 1 mM EDTA. The cell suspension was centrifuged at 500 g for 5 min at 4°C and the cell pellet was incubated with ACK lysing buffer (Thermo Fisher Scientific, A1049201) for 5 minutes to lyse the red cells. Cells were collected at 500 x g for 5 minutes then washed once in PBS. The cell pellet was resuspended in RPMI medium (Thermo Fisher Scientific, 11875-093) supplemented with 10% FBS (Gibco, 10437-028), 1% streptomycin/penicillin (Gibco, 15140-122), and 1 mM sodium pyruvate (Thermo Fisher Scientific, 11360070). Macrophages were cultured at 1.6 × 10^5^ cells/well in 12-well plates. After incubation for 2 hours, cells were washed twice with PBS and media was replenished. Cells were used for experiments after 24 hours of culture.

### Bone marrow progenitor cell isolation and bone marrow-derived macrophage (BMDM) differentiation

BMDMs preparation was performed as previously described (83). L929 conditioned media which contains the macrophage growth factor M-CSF, was prepared by culturing L929 cells (ATCC) in complete DMEM (Thermo Fisher Scientific, MT10013CV) supplemented with 10% FBS, and 1% penicillin and streptomycin for 10 days at 37°C, 5% CO_2_. The L929 conditioned media, was collected, filtered (Vacuum Filter/Storage Bottle System, Corning, 431153), and stored at −80 °C until required. For isolation of BMDMs, tibias and femurs were removed from both male and female mice and flushed with media using a 26-gauge needle. Bone marrow was collected at 500 x g for 2 min at 4 °C, resuspended with complete DMEM medium and filtered through a 70-μm cell strainer (VWR international, 10199-657). Bone marrow progenitor cells were cultured in 100 mm dishes for 6-7 days in 70% complete DMEM medium and 30% L929-conditioned medium. Fresh medium (5 mL) was added on day 3. BMDMs were collected by scraping in cold PBS containing EDTA (1 mM). After centrifugation, BMDMs were seeded into 12-well plates at a density of 1.6 × 10^5^ cells/well in DMEM and incubated overnight before use.

### Human peripheral blood mononuclear cells (PBMC) isolation

PBMC isolation was performed as previously described (84). Whole blood (20 mL) was layered upon a 15 mL Ficoll-Paque cushion (GE Healthcare, 17-1440-02) in 50 mL conical tubes, and centrifuged for 40 min at 400 x g at room temperature with the brake off. The layer of mononuclear cells at the plasma-density gradient medium interface was transferred to a new 50 mL conical tube, and 1x PBS was added to 45 ml total volume. Cells were centrifuged for 10 min at 2000 x g at room temperature and the remaining pellet of mononuclear cells was resuspended in complete RPMI medium at 1.6 × 10^5^ cells/well in 12-well plates for future experiments.

### THP-1 macrophage differentiation

Human THP-1 monocytes were differentiated into macrophages by 24 hr incubation with 100 nM PMA (Sigma-Aldrich) in complete RPMI medium at 1.6 × 10^5^ cells/well in 12-well plates. Cells were washed twice with 1x PBS and incubated with complete RPMI medium without PMA for 24 hr before experiment.

### ASC-GFP-overexpressed THP-1 macrophage

A lentivirus expressing the ASC-GFP fusion protein was prepared by transfection of pLEX-MCS-ASC-GFP (Addgene 73957), psPAX2, and p MD.2G using Fugene 6 (Promega, PAE2693) in HEK293T cells. Culture medium was replaced after 24hr with fresh medium containing 5% FBS After a further 48 hrs virus containing medium was collected and stored in -80°C. THP-1 monocytes at 1.6 × 10^5^ cells/well on 12-well plates were infected with 400uL virus with polybrene (4 µg/ml) and centrifuged at 2500 rpm at 20°C for 90min. Fresh complete RPMI medium with 100 nM PMA was added and followed by 24 hr incubation. Cells were washed twice with 1x PBS and incubated with complete RPMI medium without PMA for 24 hr before experiment.

### Overexpressed GSDMD-NT HEK293T cells

FLAG-GSDMD-NT (80951) and FLAG-GSDMD-NT-C192A (133891) were obtained from Addgene. HEK293T cells were cultured in complete DMEM medium containing 10% FBS and 1% Penicillin/Streptomycin. Transient transfection of HEK293T cells was performed using Fugene6 (Promega) according to the manufacturer’s instructions.

### HEK293As overexpressed with NLRP3, NEK7, and ASC-GFP

HEK293A cells stably expressing mouse NLRP3 (Addgene 75127), and ASC-GFP (Addgene 73957) were established under G418 (1mg/ml) and puromycin (1µg/ml) selection following transfection with Fugene6. NEK7 (Addgene 75142) was expressed in the stable cell line by transient transfection using Fugene6 for 48 hours. Cell extracts were prepared after LPS/Nigericin treatment with or without HNE for 30 min.

### LPS stimulation

Macrophage cultures were rinsed twice with serum and antibiotic-free medium (RPMI 1640 for peritoneal macrophages, THP-1 macrophages, and PBMCs or DMEM medium for BMDMs). Cells were exposed to LPS (100 ng/mL) for 3 hr with ethanol or HNE (0.3-3 µM, Millipore Sigma, 393204) in appropriate growth media. For time-course studies, individual wells were exposed to LPS (100 ng/mL) for different times, but to minimize manipulation of each plate, the total incubation time was 3 hours for all treatments.

### Inflammasome stimulation

To activate the NLRP3 inflammasome, cells were primed for 3 h with LPS (100 ng/mL) followed by addition of 2 µM (BMDMs, peritoneal macrophages, PBMCs), 6µM (THP-1 differentiated macrophages) nigericin (Sigma-Aldrich, N7143-5MG), 2 mM (peritoneal macrophages) ATP or 50 µM (THP-1 differentiated macrophages) R837 (InvivoGen, tlrl-imqs) for 30-60 min. To activate the AIM2 inflammasome, THP-1 macrophages or peritoneal macrophages were primed with LPS (100 ng/mL) for 3 hr followed by (Poly(dA:dT), 2 µg/mL) for 6 hr using LyoVec^TM^ (InvivoGen, tlrl-patc) according to the manufacturer’s protocol. To activate the NLRC4 inflammasome, BMDMs were primed with LPS (100 ng/mL) for 3 hr followed by (flagellin isolated from P. aeruginosa, 2 or 5 µg/mL) for 3 hr using FLA-PA Ultrapure (InvivoGen, tlrl-pafla) according to the manufacturer’s protocol. To activate the non-canonical inflammasome, peritoneal macrophages were stimulated with LPS (100 ng/mL) for 3 hr followed by intracellular LPS delivery (1 µg/mL) by transfection using lipofectamine 2000 (Invitrogen, 11668027), according to the manufacturer’s protocol, in OptiMEM (Gibco, 31985-070) for 16hr. HNE (0.3-10 µM) (Millipore Sigma, 393204), alkyne-HNE (1-20 µM) (Cayman Chemical, 13265) or HNE-GSH (3-10 µM) (Cayman Chemical Company, 10627) was added 30 min before the inflammasome activator.

### Cell morphology

Micrographs of cell cultures were obtained under phase contrast illumination using 40X objective (Leica) prior to cell lysis. Multiple random fields were captured for each well.

### Cell death LDH assay

Culture supernatants were collected and centrifuged at 500 × g for 5 min to remove cellular debris. LDH measurement was performed with the CyQUANT™ LDH cytotoxicity assay kit (Thermo Fisher Scientific, C20301) according to the manufacturer’s instructions. Data were plotted normalizing the O.D. value obtained in wells treated with Triton X-100 (0.1%) as 100%.

### Cell death by SYTOX^TM^ green

THP-1 macrophages or BMDMs were seeded in 96-well plates (2x10^4^ cells/well) one day before the experiments. Cells were washed twice and incubated with LPS (100 ng/mL) in XF based medium (Agilent, 103334-100) supplemented with 4.5 g/L glucose, 2 mM glutamine, 1 mM sodium pyruvate, and 1 mM HEPES buffer at final pH7.4 for 3 hr. After 3 hr LPS stimulation, SYTOX Green (final concentration 1µM) (Thermo Fisher Scientific, S7020) was added together with nigericin (2 µM for BMDMs ; 6 µM for THP-1 macrophages) and fluorescence signals (Excitation wavelength: 485 nm, Emission wavelength: 550 nm) were analyzed using FLUOstar OPTIMA plate reader (BMG Labtech) at 36 °C for 120 min. The percentage cell death was calculated by normalizing fluorescence signals from cells treated with Triton X-100 (0.1%).

### Nuclear translocation of Nrf2 and p65

Nrf2 or NF-κB p65 nuclear translocation was measured by immunofluorescence staining. THP-1 monocytes were seeded on glass coverslips in 12-well plates and differentiated as described above. Cells were treated with LPS (Nrf2 for 3 hr; p65 for 1 hr) and were washed twice with cold PBS and fixed with 4% formaldehyde in PBS for 10 min. After washing with PBS, the cells were incubated with blocking solution (10% normal goat serum, and 0.1% Triton X-100 in PBS) for 1 h followed by incubating with the Nrf2 (Abcam, ab31163) 1:500; NF-κB p65 (D14E12) (Cell Signaling Technology, 8242) 1:500; antibody overnight. Cells were washed three times with PBS and incubated with the goat anti-rabbit IgG (H+L) secondary antibody, alexa fluor 488 (Thermo Fisher Scientific, A-11034) for 1 h in PBS in the dark. After three washes with PBS, samples were mounted using fluoromount-G-DAPI (SouthernBiotech, 0100-20). Immunofluorescence was analyzed by confocal microscopy. Nuclear translocation was evaluated using NIH ImageJ software with an intensity ratio nuclei cytoplasm macro. Quantification of translocation was expressed as the percentage of intensity in nucleus over total cell (85). The data points shown are from 3 separate experiments with duplicate or triplicate wells. Around 200 cells from at least 6 images were quantified for each experimental group.

### Cytokine and HNE assays

Plasma, BAL fluid and peritoneal fluid levels of IL-1 β (Invitrogen 88-7013-22), IL-18 (Invitrogen, BMS618-3) and TNF-α (BioLegend, 430904) collected from mouse studies were determined by ELISA kits according to the manufacturer’s instructions. For in vitro studies, culture media was collected immediately after treatment. Samples were cleared by centrifugation at 16,000 ×g for 5 min and stored at −20 °C. HNE (MyBioSource.com, MBS7606509), Human and mouse IL-1 β (BioLegend, 437004 and 432604) were measured in culture supernatants by ELISA.

### RNA extraction and Real-time PCR

RNA was extracted from lung tissue or cultured cells using RNeasy kit (Qiagen, 74106) according to the manufacturer’s instructions. Complementary DNA was synthesized from 0.5 μg RNA by iScript™ cDNA Synthesis Kit (Bio-Rad, 1708891). Amplification reactions contained a target specific fraction of cDNA and 1 μM forward and reverse primers in iQ™ SYBR® Green Supermix (Bio-Rad, 1708882). Fluorescence was monitored and analyzed in a CFX connect real-time PCR system (Bio-Rad). Gene expression was normalized to β-actin using the delta delta cycle threshold method. Amplification of specific transcripts was confirmed by melting curve analysis at the end of each PCR experiment. The primers used are as follows: Mouse GCLC (Forward: AGATGATAGAACACGGGAGGAG , Reverse: TGATCCTAAAGCGATTGTTCTTC); Mouse Ferroportin-1(Forward: ACCCATCCCCATAGTCTCTGT, Reverse: ACCGTCAAATCAAAGGACCA); Mouse β-actin (Forward: TTCAACACCCCAGCCATGT, Reverse: GTAGATGGGCACAGTGTGGGT); Mouse IL-1β(Forward: GAGTGTGGATCCCAAGCAAT, Reverse: ACGGATTCCATGGTGAAGTC); Mouse IL-6 (Forward: GAGGATACCACTCCCAACAGACC, Reverse: AAGTGCATCATCGTTGTTCATACA); Mouse TNF-α (Forward: TCTTCTCATTCCTGCTTGTGG, Reverse: GGTCTGGGCCATAGAACTGA); Mouse NLRP3 (Forward: TTCCCAGACACTCATGTTGC, Reverse: AGAAGAGACCACGGCAGAAG); SAA3 (Forward: TTTCTCTTCCTGTTGTTCCCAGTC, Reverse: TCACAAGTATTTATTCAGCACATTGGGA)

### Western blot

Proteins were separated by SDS-PAGE through 10% acrylamide gels and transferred to nitrocellulose membranes, blocked with 5% nonfat dry milk in Tween-TBS and reacted with the indicated antibody: NLRP3 (AdipoGen, AG-20B-0014-C100) 1:1000; Caspase-1(AdipoGen, AG-20B-0042-C100) 1:1000; GSDMD (Abcam, ab209845) 1:1000; IL-1β (GeneTex, GTX10750) 1:2000; ASC (Santa Cruz Biotechnology, sc-514414) 1:1000; GAPDH (Millipore, MAB374) 1:2000 β-actin (Cell Signaling Technology, 4970) 1:4000; Phospho-NF-κB p65 (Ser536) (93H1) (Cell Signaling Technology, 3033S) 1:1000; IκB-α (L35A5) (Cell Signaling Technology, 4814) 1:1000; Acetyl-α-Tubulin (Lys40) (5335T) 1:1000; NEK7 (Abcam, ab133514) 1:1000 overnight. Membranes were rinsed and incubated with horseradish peroxidase conjugated secondary antibody (Anti-mouse IgG, HRP-linked Antibody (Cell Signaling Technology, 7076), Anti-rabbit IgG, HRP-linked Antibody (Cell Signaling Technology, 7074), Anti-goat IgG, HRP-linked Antibody (Jackson Immuno Research, 805-035-180). Reactive proteins were detected by enhanced chemiluminescence, visualized by exposure to radiographic film and quantified by scanning densitometry normalized to β-actin expression measured in each sample on the same gel.

### ASC speck formation

ASC-GFP-expressing THP-1differentiated macrophages were seeded on glass coverslips in 12-well plates. Cells were incubated with 100 ng/ml LPS for 3 hr followed by incubation with 6 µM nigericin for 2 h. Cells were fixed in 4% paraformaldehyde for 10 min. After three washes with 1x PBS, cells were mounted using fluoromount-G-DAPI. Immunofluorescence was analyzed by confocal microscope. The GFP 488 nm fluorescent signals were acquired by confocal microscopy. The data points represent biological replicates from 3 separate experiments. Around 200 cells from at least 6 images were quantified for each experimental group.

### ASC oligomerization

For ASC oligomer cross-linking, cells were lysed in buffer (0.5% Triton × 100, 20 mM HEPES-KOH, pH 7.5, 150 mM KCl, and complete protease and phosphatase inhibitor cocktail) on ice by syringing 10 times through a G26 needle. The cell lysates were centrifuged at 6000 rpm at 4 °C for 10 min. The pellets were resuspended in PBS and crosslinked with 2mM disuccinimidyl suberate (Thermo Fisher Scientific, 21655). The cross-linked pellets were centrifuged at 15000 rpm for 15 min and dissolved directly in a 1x SDS sample buffer.

### MitoSOX red imaging

THP-1differentiated macrophages were seeded on glass coverslips in 12-well plates. Cells were incubated with 100 ng/ml LPS for 3 hr followed by incubation with 2 µM Mitosox red (Thermo Fisher Scientific, M36008) for 30min. Cells were washed twice with medium and stimulated with nigericin for 1 hr. Cells were fixed in 4% paraformaldehyde for 10 min. After three washes with 1x PBS, cells were mounted using fluoromount-G-DAPI. Immunofluorescence was analyzed by confocal microscope. The 510/580 nm fluorescent signals were acquired by confocal microscopy.

### Click chemistry and biotin affinity precipitation

Cells were lysed in IP Lysis buffer (20 mM HEPES-KOH, 150 mM NaCl, and 0.5% v/v Triton-100x) containing protease inhibitor cocktail, and treated with 10 mM sodium borohydride (Sigma-Aldrich, 452882) for 30 min. Click-chemistry was performed with Molecular Probes Click iT Protein Reaction Buffer Kit (Thermo Fisher Scientific, C10276) according to the manufacturer’s instructions. Briefly, The Cu(I)-catalyzed click reaction was initiated by adding Click-iT reaction buffer and 50 µl of Azide-PEG3-biotin in 50 mM (Cayman Chemical, 23419) to each sample and incubated on a rotator for 20 min at room temperature. Protein was precipitated and the pellet was washed once with 100 µl of ice-cold methanol and centrifuged at 15,000 x g for 5 min. Methanol was removed and the pellets were re-suspended in 50 µl IP Lysis buffer containing protease inhibitor cocktail. Protein concentrations were estimated assuming no protein loss occurred during the click and precipitation procedure. To perform biotin affinity precipitation, clicked lysates were loaded with 20 µl of Dynabeads™ MyOne™ Streptavidin C1 (Invitrogen, 65001). The Dynabeads were equilibrated to buffer by three 100 µl washes. Thirty micrograms of “clicked” lysates were loaded onto the equilibrated beads. The volume was brought up to 100 µl with IP lysis buffer, and lysates were incubated with beads for 1 h at room temperature on a rotator. After incubation, the beads were washed six times with 100 µl volumes of IP lysis buffer. The bound proteins were eluted using 100 µl of 1X sample lysis buffer, by vortexing and heating at 95 °C for 10 min. The supernatant was collected and stored at −20 °C for western blot analysis.

### Co-immunoprecipitation

Cells were lysed in IP buffer on ice by syringing 10 times through a G26 needle. The supernatant was collected as the whole cell lysate (WCL) sample. One mg total protein from WCL was incubated with anti-NEK7 antibody (Abcam, ab133514), anti-NLRP3 antibody (AdipoGen, AG-20B-0014-C100), ChromPure Rabbit IgG (Jackson Immuno Research, 011-000-003), or ChromPure Mouse IgG (Jackson Immuno Research, 015-000-00) overnight. Agarose beads (Roche, C755B62) was added to the remaining supernatant, which was subsequently incubated at 4°C in a ferris wheel mixer for 3 hr. IP samples were subsequently centrifuged at 200 x g for 2 min at 4°C, supernatant removed, and beads washed five times with 1 mL low stringency lysis buffer. The immune complexes were eluted by addition of 100 µL of 1X sample lysis buffer, boiled for 5 min and analyzed by western blot.

### Statistics

Unless otherwise noted, in vitro experiments were repeated as three independent procedures, with duplicate or triplicate wells averaged prior to statistical analysis. All data were presented as mean ± SD. GraphPad Prism 8.0 was used for statistical analysis. Comparisons between two groups after stimulation were analyzed by two-way ANOVA. HNE dose response experiments in cell cultures were analyzed by one-way ANOVA followed by post hoc T tests using Bonferroni correction for multiple comparisons. P values were indicated as follow: * < 0.05, ** < 0.01, *** < 0.001, **** < 0.0001.

## Author Contribution Statement

C.G.H., M.S., C.Y., B.C.B. designed research; C.G.H., C.L.C., C.Z., M.S. performed research; B.C.B. contributed new reagents/ analytic tools; C.G.H., C.Z. analyzed data; C.G.H., C.Y., B.C.B. wrote the paper.

## Acknowledgments

None

## Ethics Statement

All of the experiments were approved by the University Committee on Animal Use For Research (UCAR) at the University of Rochester and followed National Institutes of Health guidelines for experimental procedures on mice. Human blood samples from healthy donors were collected and processed at the University of Rochester Medical Center following Institutional Review Board approval.

## Data Availability Statement

All data generated or analysed during this study are included in this published article and in its supplementary file.

