## Supplementary Information for "The lipid peroxidation product 4-hydroxynonenal inhibits NLRP3 inflammasome activation and macrophage pyroptosis"

**This PDF file includes:**

Supplementary Figures S1 to S13

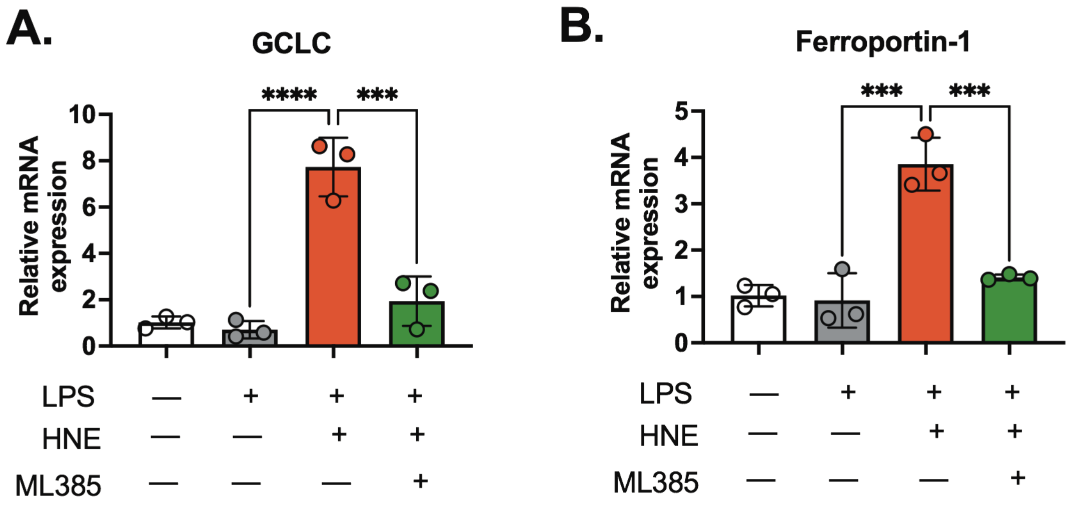


**Supplementary Figure S1. HNE-induced Nrf2 signaling was inhibited by ML385.**

**A-B.** Peritoneal macrophages were stimulated with or without LPS (100 ng/mL), HNE (3 μM) or ML385 (2 μM) for 3 hr. Gene expression was analyzed by real-time PCR. (A) GCLC mRNA expression and (B) Ferroportin-1 mRNA expression were normalized to β-actin.

Statistics in A-B were performed using an one-way ANOVA and Bonferroni's post hoc test. $P<0.05$, among treatment groups. Bars represent mean ± SD.

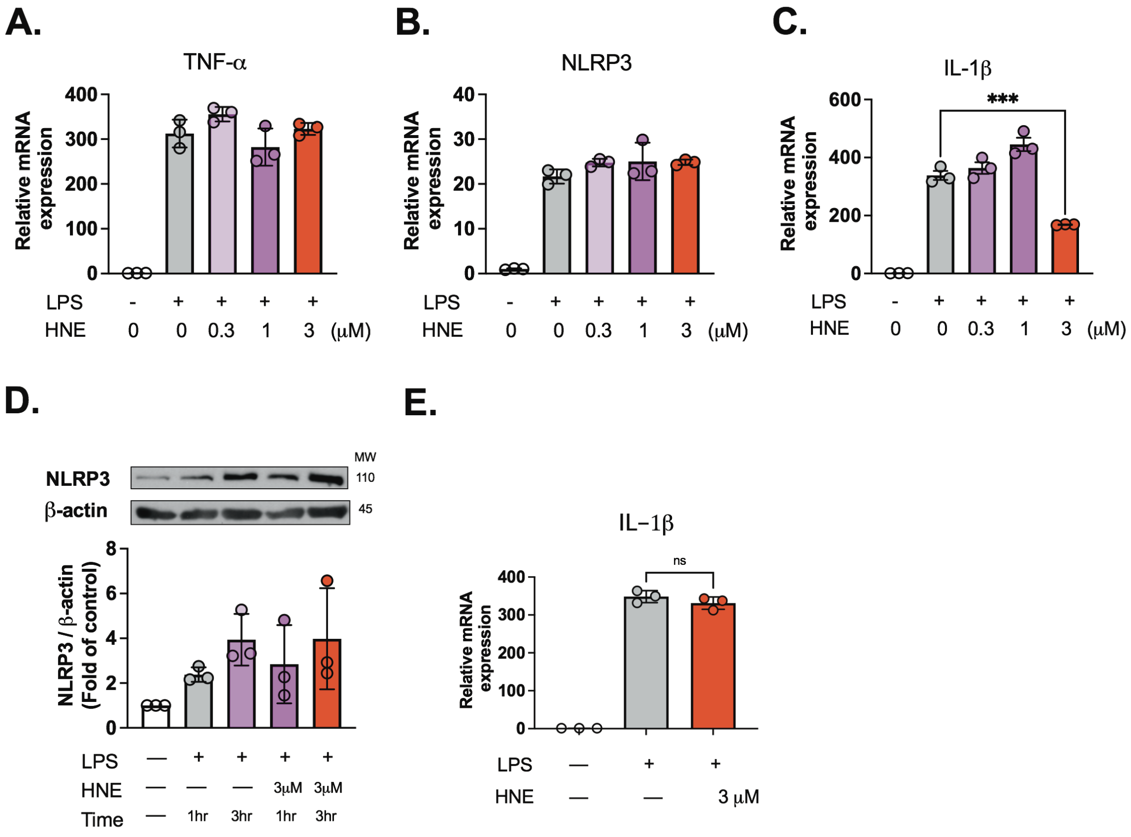


**Supplementary Figure S2. HNE does not inhibit NLRP3 and TNF-α transcriptional responses to LPS stimulation in mouse macrophages.**

**A-C.** Peritoneal macrophages were stimulated with LPS (100 ng/mL) and co-incubated with ethanol or HNE (0.3-3 μM) for 3 hr. Cell lysates were analyzed by real-time PCR. (A) TNF-α and (B) NLRP3 (C) IL-1β. N=3 independent experiments.

**D.** Peritoneal macrophages were stimulated with LPS (100 ng/mL) and co-incubated with ethanol or HNE (3 μM) for 1or 3 hr. Cell lysates were analyzed by western blot. (D) NLRP3, and β-actin western blots are representative of three independent experiments.

**E.** Peritoneal macrophages were stimulated with LPS (100 ng/mL) for 3 hr. HNE (3 μM) or ethanol was added 30 min before harvesting. Gene expression was analyzed by real-time PCR. (E) IL-1β. N=3 independent experiments.

Statistics in A-E were performed using a one-way ANOVA and Bonferroni's post hoc test. $P<0.05$, among treatment groups. Bars represent mean ± SD.

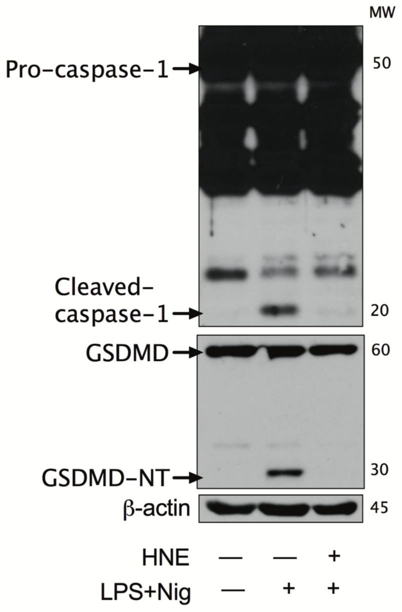


**Supplementary Figure S3. HNE inhibits NLRP3 inflammasome via a transcription-independent mechanism.**

BMDMs were pre-treated with HNE for 30 min, and then stimulated with nigericin (Nig, 2 μM) after 10 min LPS (100 ng/mL) priming. Western blots are representative of three independent experiments.

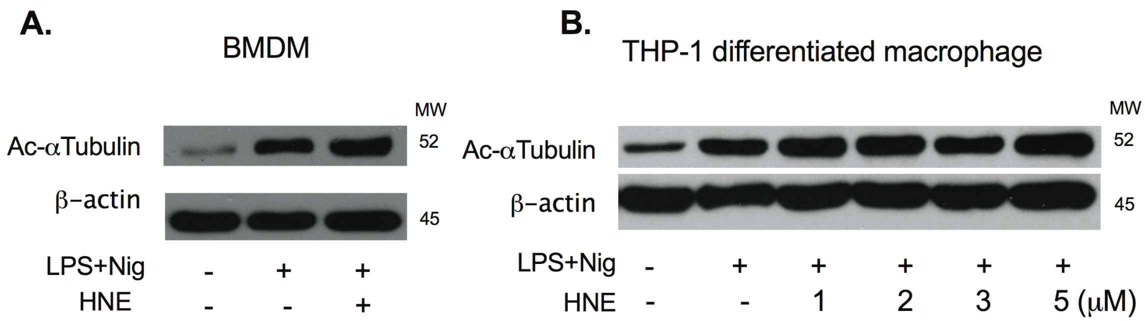


**Supplementary Figure S4. HNE has no effect on nigericin-stimulated α-Tubulin acetylation in LPS primed macrophages.**

**A-B.** (A) BMDMs (Bone marrow-derived macrophages) and (B) THP-1 differentiated macrophages were stimulated with LPS (100 ng/mL) for 3 hr followed by nigericin (Nig, 2 μM). HNE (3 μM or indicated concentration) was added 30 min before nigericin. Acetyl-α-Tubulin (Lys40) western blots are representative of three independent experiments.

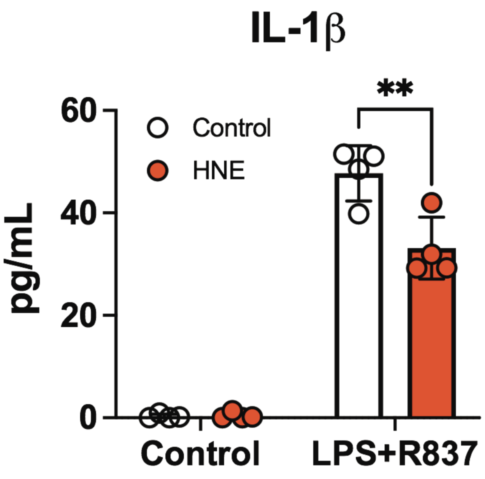


**Supplementary Figure S5. HNE inhibits non-potassium efflux-mediated NLRP3 activation.** THP-1 macrophages were stimulated with LPS for 3 hr followed by 100 µM R837 for 1 hr. Ethanol or HNE (3 µM) was co-incubated with cells 30 min before R837 (50 µM). IL-1β in the medium was measured by ELISA.

Statistics were performed using a 2-way ANOVA and Bonferroni's post hoc test. **P<0.01 between control and treatment groups. Bars represent mean ± SD.

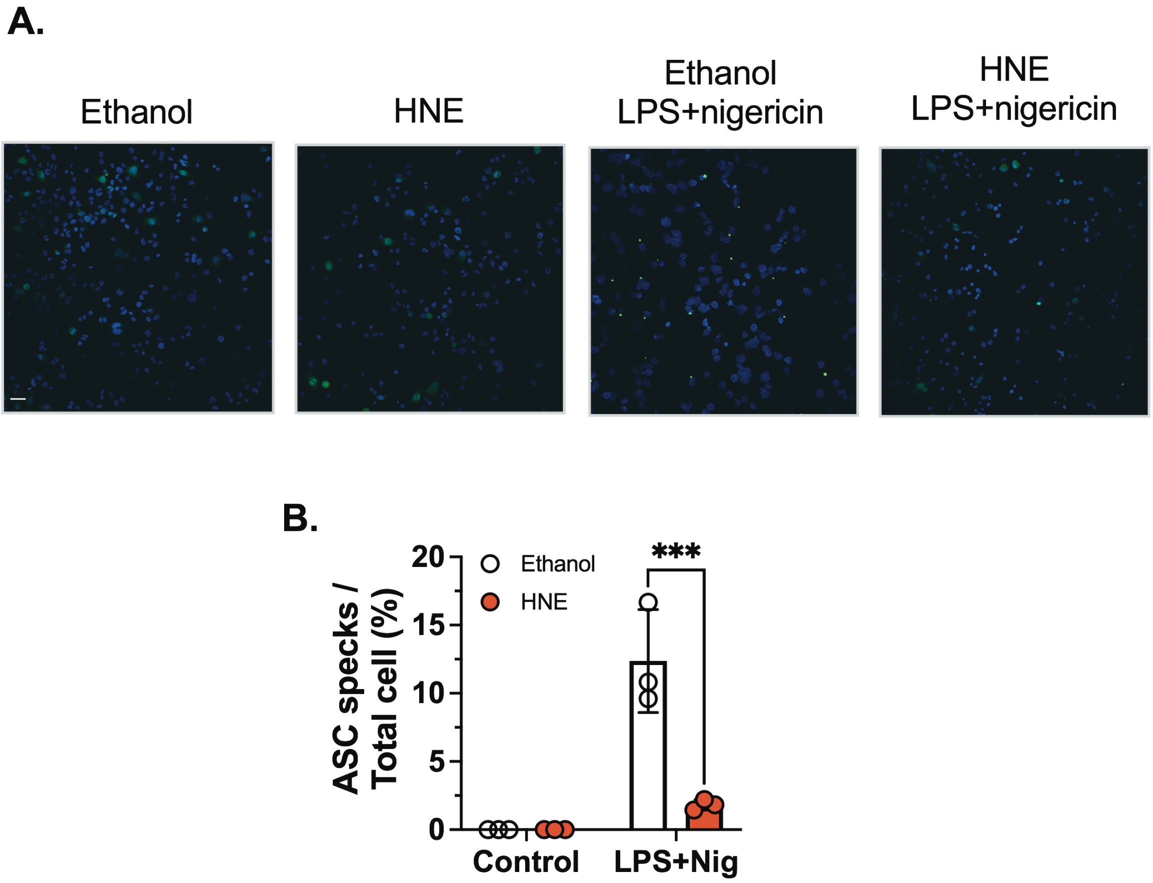


**Supplementary Figure S6. HNE inhibits ASC speck formation.** THP-1 macrophages that overexpressed ASC-GFP were stimulated with LPS (100 ng/mL) followed by nigericin (Nig, 6 μM) for 2 hr. ASC speck formation (green) was measured by confocal microscopy. (scale bar: 50 μm ), and quantified by Image J.

Statistics in B were performed using a 2-way ANOVA and Bonferroni's post hoc test. ***$P<0.001$ between control and treatment groups. Bars represent mean ± SD.

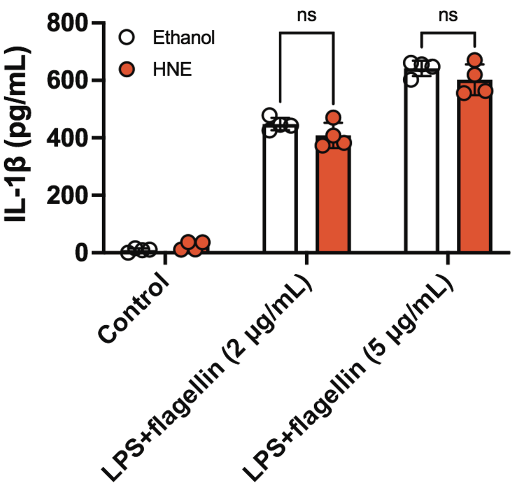


**Supplementary Figure S7. HNE has no effect on NLRC4 activation.**

BMDMS were stimulated with LPS (100 ng/mL) for 3 hr followed by flagellin transfection (2 or 5 μg/mL) for 3 hr. Ethanol or HNE (3 μM) was co-incubated with cells 30 min before transfection. IL-1β was measured by ELISA.

Statistics were performed using a 2-way ANOVA and Bonferroni's post hoc test. $P<0.05$ between control and treatment groups. Bars represent mean ± SD.

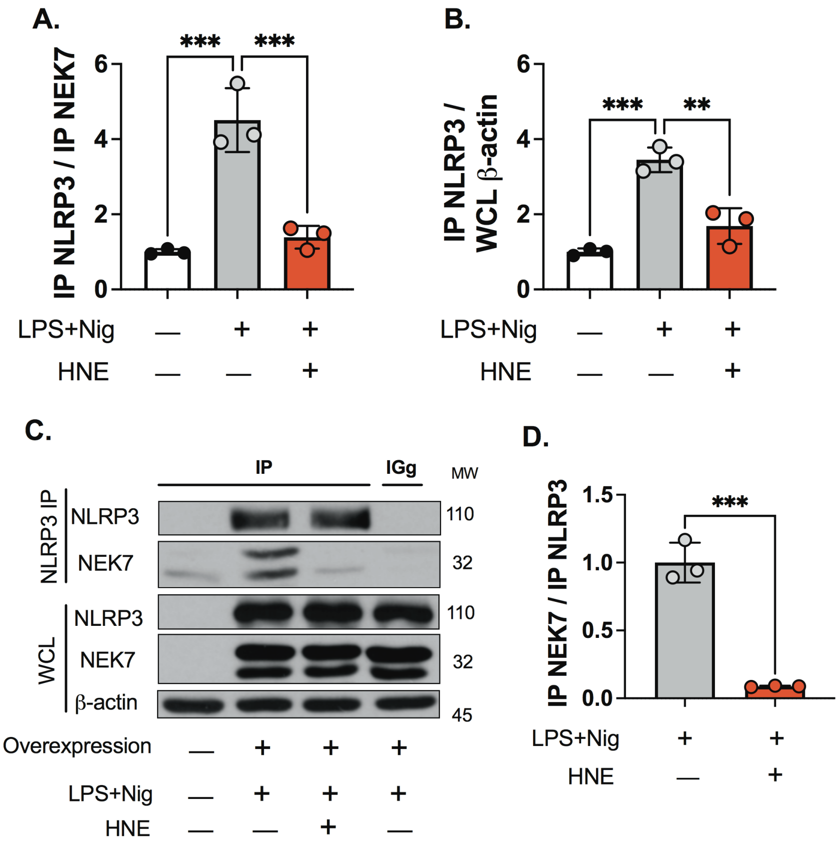


**Supplementary Figure S8. HNE blocks the interaction between NLRP3 and NEK7**

**A-B**. BMDM were stimulated with LPS (100 ng/mL) for 3 hr followed by nigericin (Nig, 2 μM) for 60 min. Ethanol or HNE (3 μM) was added 30 min before nigericin. Immunoblots of NLRP3, NEK7, and β-actin from NEK7-immunoprecipitated and whole cell lysates (WCL) are quantified by three independent experiments.

**C-D.** Stable HEK293A cell lines overexpressing NLRP3, NEK7, and ASC-GFP NLRP3 were stimulated with LPS (100 ng/mL) and nigericin (6 μM) for 30min. (C) Representative western blots from 3 independent experiments (D) The quantification data were from 3 different experiments.

Statistics in A, B and D were performed using an one-way ANOVA and Bonferroni's post hoc test. ***$P<0.001$ , **$P<0.01$ between control and treatment groups. Bars represent mean ± SD.

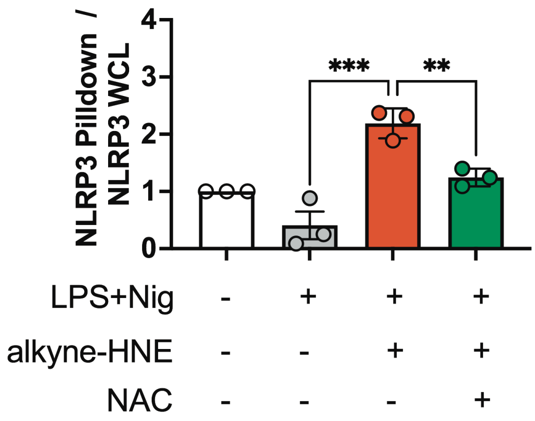


**Supplementary Figure S9. Alkyne-HNE pulls down NLRP3.**

BMDM were stimulated with LPS (100 ng/mL) for 3 hr followed by nigericin (Nig, 2 μM) for 30 min. Alkyne-HNE (10 μM), N-acetyl cysteine (NAC, 500 μM), or both were added 30 min before nigericin. Immunoblots of NLRP3 from “clicked” and whole cell lysates (WCL) are quantified from 3 independent experiments.

Statistics were performed using an one-way ANOVA and Bonferroni’s post hoc test. ***P<0.001 , **P<0.01 between control and treatment groups. Bars represent mean ± SD.

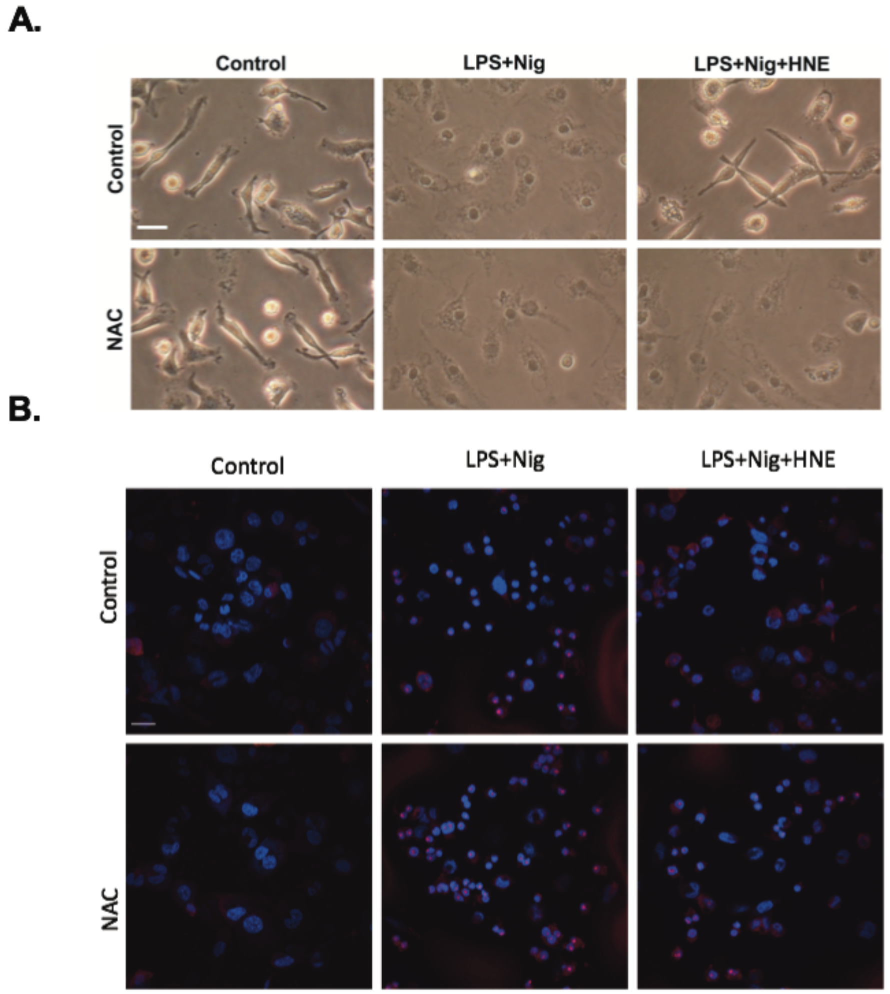


**Supplementary Figure S10. HNE decreases pyroptosis and MitoSOX accumulation in the nuclear area after LPS+nigericin stimulation through a cysteine-dependent mechanism . A.** Peritoneal macrophages were stimulated with LPS (100 ng/mL) for 3 hr followed by nigericin (Nig, 2 μM) for 1 hr. HNE (3 μM), n-acetyl cysteine (NAC, 500 μM), or both were co-incubated with cells. Representative images of cell cultures were obtained under phase contrast illumination using 40X objective (Leica). (scale bar: 10 μm ).

**B.** THP-1 macrophages were stimulated with LPS (100 ng/mL) for 3 hr followed by nigericin (Nig, 2 μM) for 1 hr. HNE (3 μM), n-acetyl cysteine (NAC, 500 μM), or both were co-incubated with cells. Representative images of MitoSOX red fluorescence were measured by confocal microscopy. (scale bar: 20 μm ).

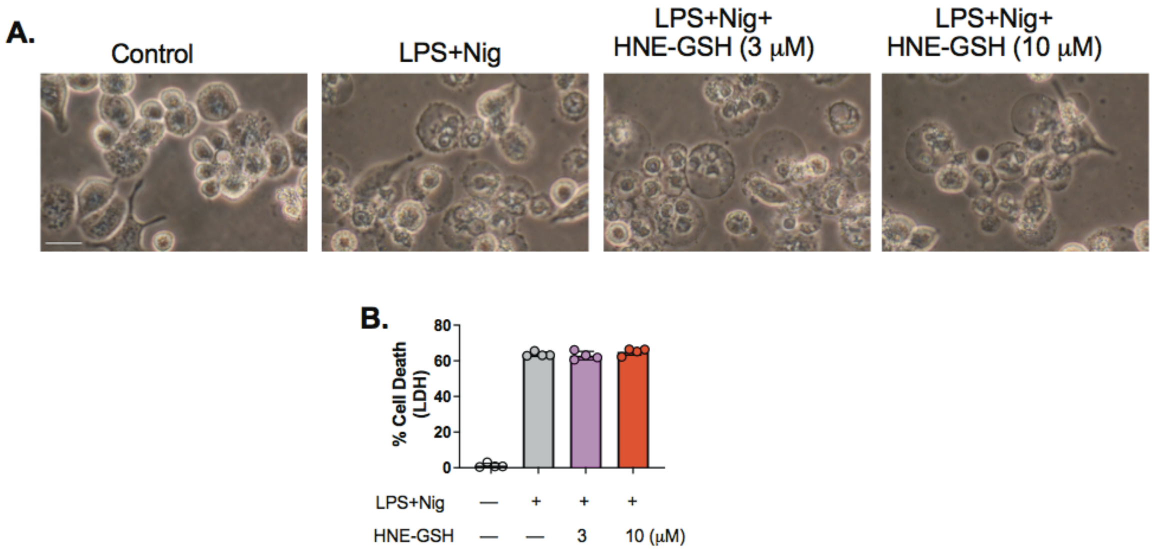


**Supplementary Figure S11. HNE-GSH has no effect on cell death after LPS+nigericin stimulation.**

**A.** THP-1 differentiated macrophages were stimulated with LPS (100 ng/mL) and co-incubated with HNE-GSH (3 or 10 μM) for 3 hr followed by 2 hr nigericin (Nig, 6 μM) treatment. (A) Morphology, scale bar = 10 μm. (B) LDH cytotoxicity.

Statistics in B were performed using an one-way ANOVA and Bonferroni's post hoc test. $P<0.05$ among groups. (N=4 experiments). Bars represent mean ± SD.

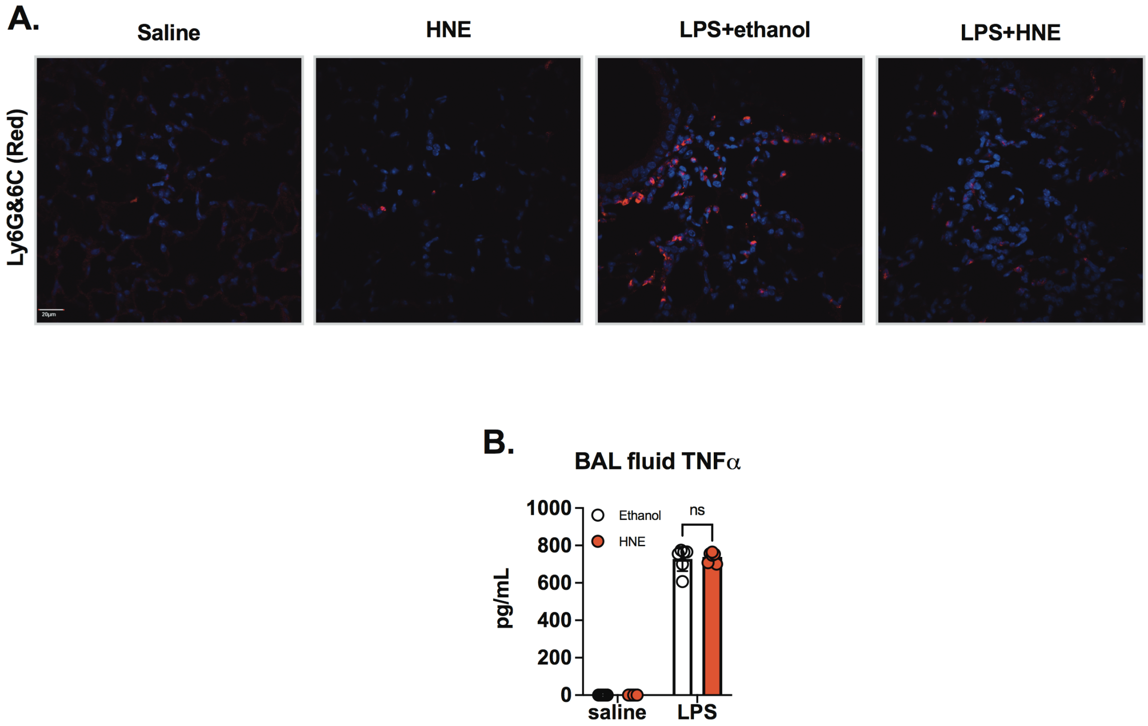


**Supplementary Figure S12. HNE inhibits neutrophil and macrophage infiltration in mouse acute lung injury model.**

**A-B.** Ethanol (0.01%) in saline (50 µl), HNE (6 µM) in saline (50 µl), LPS (2 mg/kg) + ethanol (0.01%) in saline, or LPS (2 mg/kg) + HNE (6 µM) in saline were delivered oropharyngeally to mice. Lung tissues were harvested at 3 hr post treatment for immunohistochemistry (N=4 mice from each group). BAL fluid was harvested at 18 hr (N=4 mice from each saline group, and N=6 from each LPS groups). (A) Immunofluorescence of representative lung sections for Ly6G&6C  (Red), DAPI (Blue). Quantification results of (B) TNF-α protein in BAL fluid.

Statistics were performed using a 2-way ANOVA and Bonferroni's post hoc test.  $P<0.05$ between control and treatment groups. Bars represent mean ± SD.

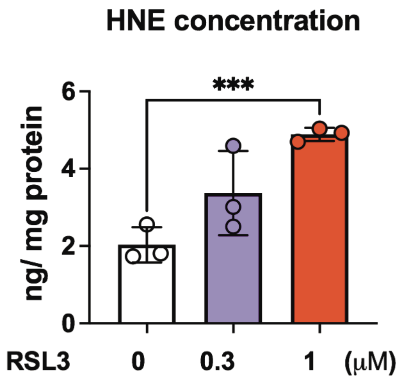


**Supplementary Figure S13. RSL3 increases HNE in a dose-dependent manner.**

**A.** BMDMs were stimulated with LPS (100 ng/mL) and co-incubated with RSL3 (3 or 10 μM) for 3 hr (A) HNE levels were measured by ELISA.

Statistics were performed using an one-way ANOVA and Bonferroni's post hoc test. ***P<0.001 among groups. (N=3 experiments). Bars represent mean ± SD.
